# The Art of Brainwaves: A Survey on Event-Related Potential Visualization Practices

**DOI:** 10.1101/2023.12.20.572507

**Authors:** Vladimir Mikheev, René Skukies, Benedikt V. Ehinger

## Abstract

Electroencephalography (EEG) and event-related potentials (ERPs) have been analyzed for more than 70 years. Yet, we know little about how practitioners visualize the results of their analyses. Here, we designed an online survey (n=213) targeting M/EEG practitioners from novice to expert level. Our primary goal is to better understand the visualization tools currently in use, the challenges researchers face, and their experiences and opinions on how best to display their brain data. Finally, we explored whether researchers are aware of more general visualization issues. In this paper we provide an overview of the most popular ERP visualization tools. Then, we found that the community does not have a unique nomenclature to refer to some plot types, and we propose a set of recommendations to name the most popular ERP plot types. Finally, we provide an analysis of practitioner feature preferences for software developers and conclude with further recommendations for ERP practitioners.

## 1. Introduction

Electroencephalography (EEG) is used to detect and record electrical voltages of the brain. These voltages are recorded through an array of electrodes placed on the scalp, resulting in a multivariate timeseries. For analysis, the data are typically transformed into event-related potentials (ERPs) by aligning the timeseries to events of interest and averaging over repeated events per condition. Commonly, the results consist of an array with four dimensions: sensors x time x conditions x subject (Figure 1).

**Figure 1.**
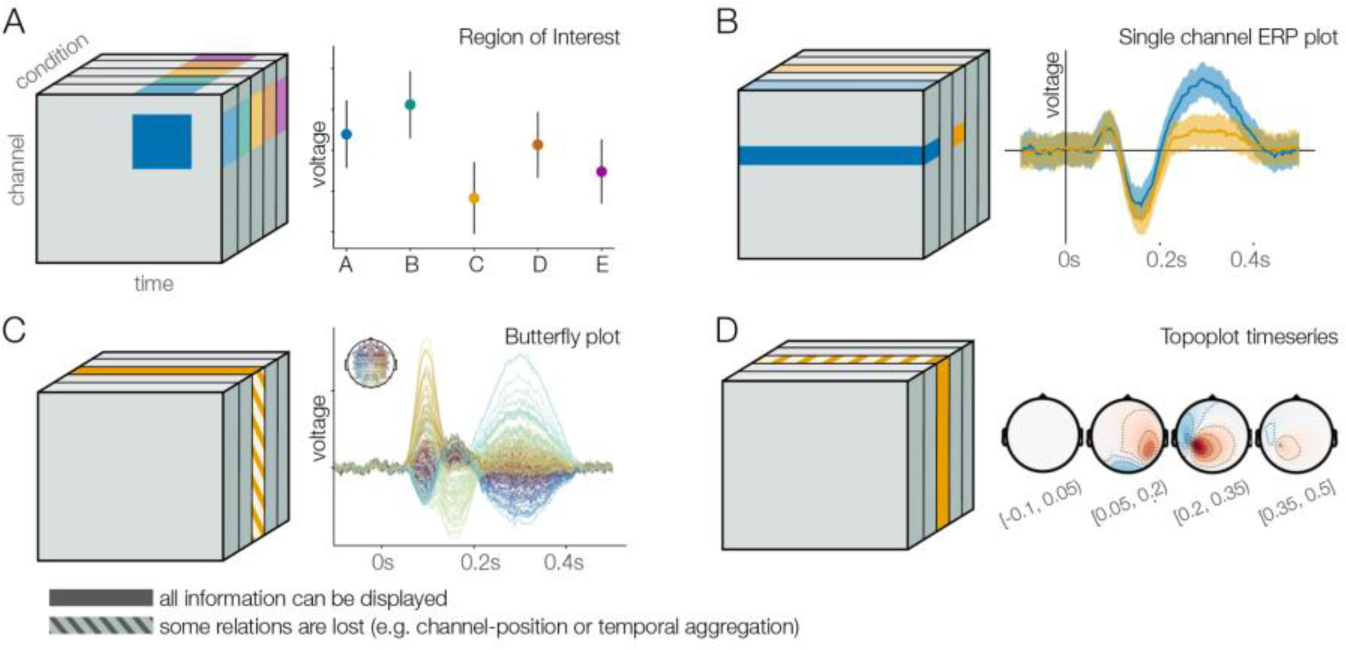
ERP dimension cube. It is possible to visualize some aspects of a dataset by averaging in time and space (A), sub-selecting channels and conditions (B), or omitting some information of a dimension (C, D). Categorical color (blue, green, yellow, orange, red) indicates experimental condition.

These dimensions and their different features can be visualized using a variety of different plot types. For example, the ERP plot allows us to highlight slices of the condition and time dimension, at the cost of collapsing over the sensor and subject dimensions (Figure 1B). Contrary, if we want to display all sensors, we might want to switch to a butterfly plot (Figure 1C), a topoplot (Figure 1D), or a channel plot, at the cost of selecting a single condition. If we limit ourselves to a selection (and/or averaging) across sensors and time, we can even use more traditional scatter plots (Figure 1A). However, there is no single way to visualize all the information in this hypercube at once.

The problem of dimensionality is even more pronounced in the more general regression-based ERP framework (Smith & Kutas, 2015), which allows for the analysis of more naturalistic experiments including, eye movements (Dimigen & Ehinger, 2021) or mobile/VR EEG (Ehinger et al., 2014; Jacobsen et al., 2022; Scanlon et al., 2023; Wunderlich et al., 2023; Zink et al., 2016). In such analyses, the “condition” dimension is generalized to a “regression” dimension containing coefficients of the multiple regression, which leads to additional visualization complexities of multiple regression (Klemelä, 2014).

To emphasize a particular dimension, EEG practitioners should be aware of the available and appropriate plot types. Currently, there are plenty of ways to visualize plot types: an initial literature review revealed more than 30 different EEG analysis tools with widely varying popularity. In opposition to that, the literature on EEG visualization is scarce, so understanding of how to present EEG data properly is limited. In addition, each tool has its own approach to usability and naming of EEG visualizations, which might confuse an inexperienced user. Ideally, researchers should know what plot types exist, what kind of dimensions they can represent, how to implement a plot in a certain tool, what kind of information is crucial to show on a plot for better generalizability. And, of course, there should be a common notation to avoid any confusion.

To understand the current state of ERP visualization, we surveyed 213 practitioners, including novices and experts. We wanted to assess the challenges analysts face when visualizing EEG data, how they would like to see current tools improved, and what they know about ERP plot types, their characteristics, and their implementation.

In summary, this leads us to the following research questions:

- RQ 1: Is there a general naming convention of ERP plot types?
- RQ 2: Which ERP visualization tools are known and in use?
- RQ 3: What features are important in current ERP visualization tools and what are potential improvements for tools?
- RQ 4: What does the EEG community think about some of the controversies surrounding ERP visualization?
- RQ 5: How well are ERP researchers aware of some of the perceptual issues in visualization?
- RQ 6: Does proficiency in EEG correlate with opinions towards ERP visualizations requirements?

We believe that answering these questions will be helpful to create a better theoretical framework of ERP plot types. Such information would be helpful to improve existing visualization tools and to create new ones.

## 2. Background

In the following sections, we provide an overview of current ERP visualization types, controversies in ERP visualization, and some of the perceptual issues in visualization that directly apply to ERP researchers.

### 2.1. State-of-the-art techniques for ERP visualization

The literature on ERP visualization types is scarce. To our knowledge, only ten Caat (ten Caat, 2008; ten Caat et al., 2007) explicitly worked on this topic. They identified seven common types of EEG plots and counted the number of time steps and number channels they can display. They further focused on the use of parallel coordinates to visualize ERP results and proposed a new type of ERP visualization: tiled parallel coordinates, a multiple miniature version of parallel coordinates plots. Because this more advanced plot type has not gained traction in the field, we included only the basic parallel coordinates plot visualization in our survey (see Figure 2H).

**Figure 2.**
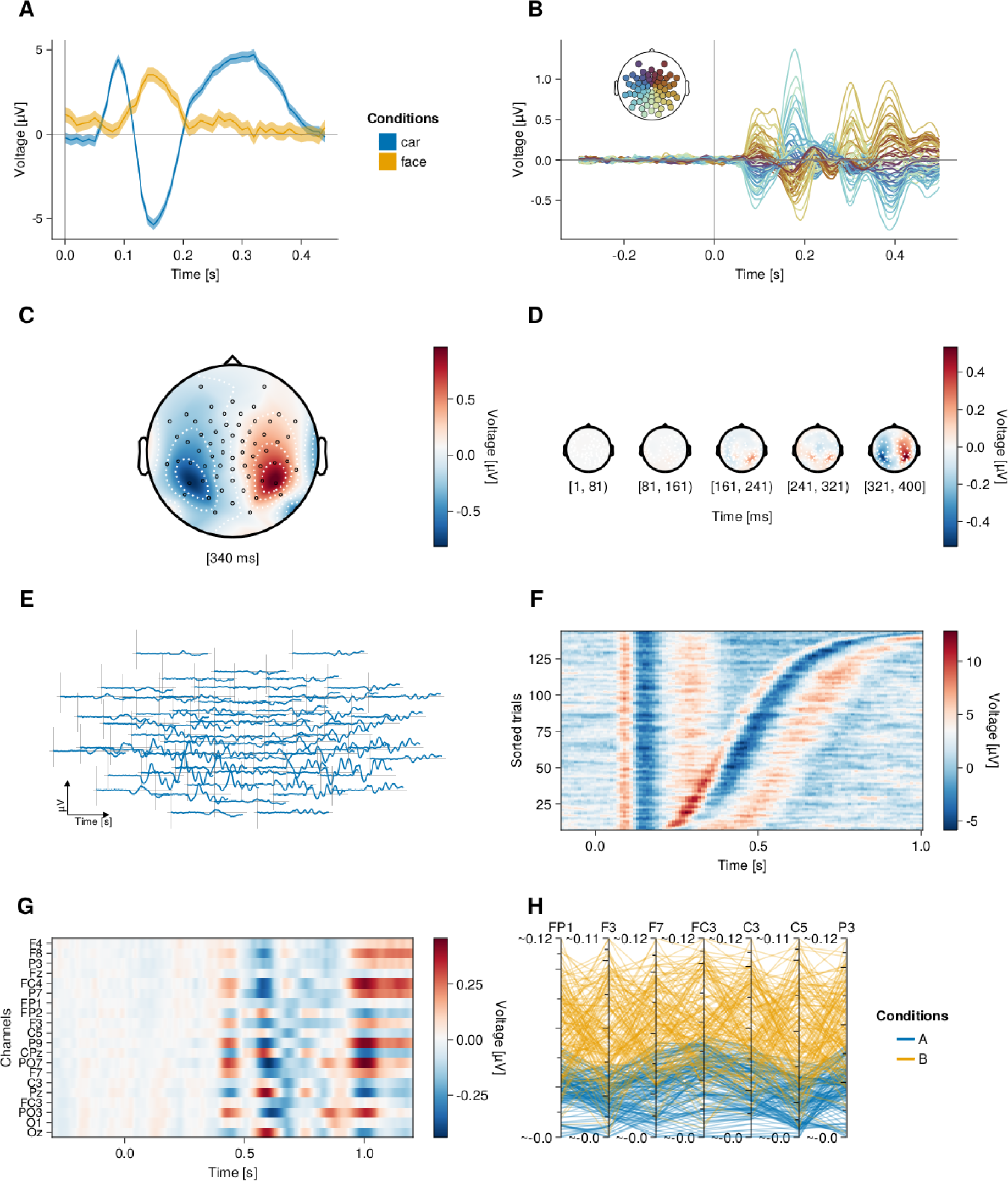
Plots of the eight visualizations used in our survey using *UnfoldMakie.jl*. A) ERP plot, B) Butterfly plot, C) Topoplot, D) Topoplot timeseries, E) ERP image, G) Channel image. H) parallel coordinate plot.

In our study, we used the Ten Caat’s list of plot types and added the channel image plot^1^ (Table 1). Note, that due to lack of adoption in the field^2^, we changed some of their suggested plot type names based on two principles. Either they were the result of a user consensus shown in this survey, or we proposed a new convention after carefully analyzing the raw responses and taking other considerations into account (see Section 2 in Discussion). Exemplary plots for all eight visualizations are shown in Figure 1. Similar plots were used in the survey (Supplementary Figure 1).

**Table 1.**
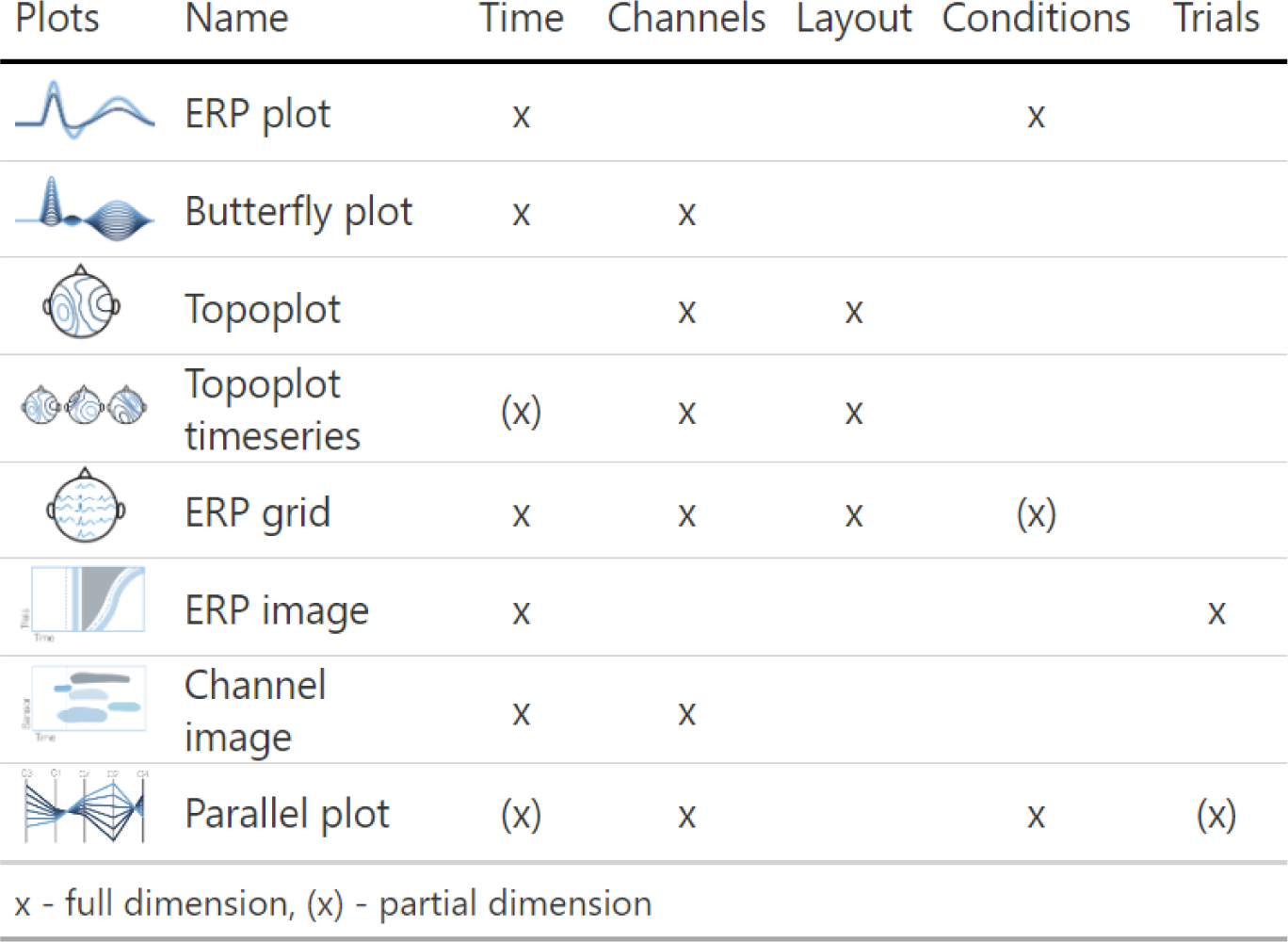
The eight ERP visualizations used in this study, and the data dimension they can effectively depict.

| Plots | Name | Time | Channels | Layout | Conditions | Trials |
| --- | --- | --- | --- | --- | --- | --- |
|  | ERP plot | x |  |  | x |  |
|  | Butterfly plot | x | x |  |  |  |
|  | Topoplot |  | x | x |  |  |
|  | Topoplot timeseries | (x) | x | x |  |  |
|  | ERP grid | x | x | x | (x) |  |
|  | ERP image | x |  |  |  | x |
|  | Channel image | x | x |  |  |  |
|  | Parallel plot | (x) | x |  | x | (x) |
| x - full dimension, (x) - partial dimension |  |  |  |  |  |  |

Previously, two initiatives proposed recommendations for plotting and categorizing ERP visualizations. First, Pernet et al. have developed COBIDAS (Committee on Best Practices in Data Analysis and Sharing), the M/EEG best practices guide (Pernet et al., 2020). Part of their recommendations relate to plotting: label all axes and report units, plot all channels and conditions, show error measures (CI or SD). Second, ARTEM-IS (Agreed Reporting Template for Electroencephalography Methodology) is a documentation tool for EEG studies (Šoškić et al., 2023; Styles et al., 2021). It facilitates reporting critical parameters of EEG analysis pipelines on replicability. Currently, it consists of 8 sections, including one for visualization. One can specify plot type and post-processing for visualization purposes.

However, while such initiatives are a step in the right direction, an earlier study showed that the publication of guidelines is not enough to change the actual methodological practices (Clayson et al., 2019). Concurrently, it could be helpful to directly implement such guidelines into the analysis software. Further, the categorisateeion of ERP plot types still hasn’t been a focus in these guidelines, which we aim to address in this paper.

### 2.2. Controversies in ERP visualization

Based on discussions, a literature review, and our own experiences, we decided to investigate several controversies in the ERP community regarding the analysis and visualization of ERP data. While calling them “controversies’’ may be a bit of a stretch, strong opinions on these issues have been expressed in private conversations and at conferences. In particular, we identified four controversies: 1) Which channels should be recorded, analyzed and visualized (single channel, region of interest, or all channels?) 2) How much, if any, of the baseline period should be visualized? 3) Should ERP positivity be plotted upwards or downwards? 4) Should we depict single timesteps or average time windows in a topoplot timeseries?

#### 1) Channel exclusion

It is common that not all measured electrodes are actually analyzed, begging the question of why they were recorded in the first place. If one studies a well-known effect, with a consensus on the timing and location, focusing on the singular necessary electrodes is best practice. In this case, only those electrodes would need to be recorded. The situation is quite different if mass-univariate analyses are required, or if “excess” electrodes are used for average referencing, independent component analysis or other multivariate preprocessing methods.

Other reasons are less well motivated, for instance if fewer channels are used solely to simplify the interpretation or to ease computational burden. In these cases, important effects might be dismissed too early. And finally, a reason relevant to this study and potentially addressable reason would be, if channels are excluded because the required visualizations are not known by users or are not readily available in software.

To shed light on the prevalence of analyzing subsets of electrodes, we included a question in our survey asking about the number of electrodes recorded and subsequently analyzed. We did not ask more detailed questions about individual researchers’ reasons for possibly not using all electrodes. Future questionnaires could explore this issue in more detail.

#### 2) Baseline visualization

Baseline correction is a data preprocessing technique commonly employed in ERP analysis or visualization, mainly used to correct for unexpected offsets between conditions. These offsets might happen due to e.g. non-electrophysiological activity (muscles, sweating etc.) triggering systematic variations in data (Urbach & Kutas, 2006). Our focus is not related to the actual baseline correction and whether it is necessary (for a discussion on this topic we recommend (Alday, 2019), but rather the visualization of the baseline period. It is commonly recommended to display some part of the baseline period to visually estimate data quality and identify potential confounds in randomization (Luck 2014). However, how much baseline period should be shown (if any) remains unclear, and no empirical data on this issue has been collected.

#### 3) Polarity convention

Contrary to the established mathematical standard, it is quite common in ERP visualization to plot negative voltages upward on the Cartesian coordinate system (see Luck 2014 Box 1.3). Defendants of this practice typically point towards historical reasons, often quoting the earliest (Davis et al., 1939) or outstanding ERP studies (Sutton et al., 1965; Walter et al., 1964). Further, some researcher explain their preference by relating ERPs to the underlying physiological processes, which are often associated with negative electrical charges (Picton, 2010)^3^. However, there is no established consensus in the EEG community on the “right” polarity convention. In this survey, we assessed how much support currently exists for either polarity preference, and how this varies across EEG research fields.

#### 4) Topoplot timeseries interpretation

Topographic timeseries show voltage differences at different locations on the scalp over time. However, it is rarely labeled whether they represent a single point in time, or an average over a period of time which might lead to ambiguity in interpretation. In the survey, we assessed whether researchers share a common intuition.

### 2.3. Common misleading visualization techniques

Inadequate visualizations have the potential to be misleading or hinder understanding of the results. There are two issues we addressed in our survey: 1) the visualization of uncertainty and 2) the use of perceptually problematic color maps.

1. Omitting uncertainty on a graph, or wrongly inferring what type of uncertainty is depicted can lead to misinterpretation of data. However, omitting uncertainty depictions is quite common in scientific papers. A study of visualizations in the more general neuroscience literature analyzed 1,451 figures from 288 articles (Allen et al., 2012). Around 20% of the 2D plots (bar/box/violin plots) lacked uncertainty labels. Of those that had uncertainty depictions, 30% did not label the type of uncertainty used (e.g., standard error bars or confidence intervals). Results for 3D graphs (i.e. heatmaps) were much worse, with 80% lacking uncertainty depictions and no uncertainty labeling in the remaining. We expect this to be even worse for time series ERP data because error bands are difficult to add to time series in most plotting tools. In our survey, we investigated how commonly uncertainty estimates are reported for time-resolved ERP plots, and what type of uncertainty is typically reported (standard error, confidence interval, or other).
2. Color maps are used to map numerical values to color, most commonly in heatmaps. However, some color maps are not “veridical”, meaning they cannot truthfully display the underlying data. Visualization researchers (Crameri et al., 2020; Moreland, 2016) have identified three major problems that can occur when using color maps.

A) Violation of perceptually uniformity. The perceived difference between two colors on the color map does not correspond to their numerical representation. For example, a segment in one part of the color map represents the hues of the same color, while in another part a segment of the same length represents two different colors. B) Violation of perceptual order. The lightness and brightness increase non-linearly, for instance first comes a bright color, then dark color, then again bright. That makes the order of colors not intuitive, and readers need to constantly check the color bar. C) Unfriendliness to colorblindness. Typically, color maps become problematic when red and green have similar luminosity, making them indistinguishable to colorblind people. Proportions of colorblind people can vary by ethnicity and gender, for example, red-green color deficiency is prevalent in ∼8% of males and 0.5% of females in the European Caucasian population (Birch, 2012).

The “jet” color map has all three problems. Which leads to the fact that the jet color map creates spurious “bands” when a uniform area is plotted (Figure 3, Quinan et al., 2019). Still, it is used as the default option in EEGLAB, the most popular EEG analysis toolbox. On this issue, we set out to identify if the perceptual controversies around the color map (by way of example of the “jet” color map) are known to researchers.

**Figure 3.**
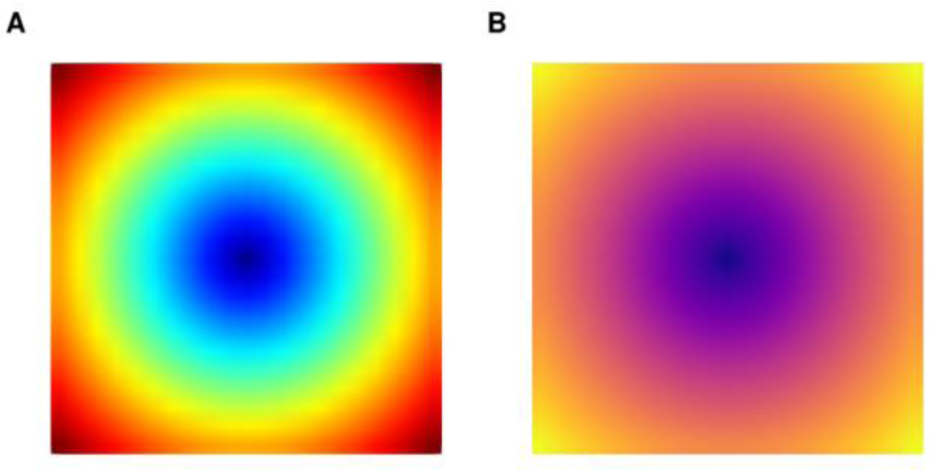
Example of two-color maps depicted as a linear center-surround gradient: Jet (A) and Plasma (B) (Smith et al., 2015). We can see that Jet creates spurious “bands” while Plasma color map shows a linear change.

## 3. Methods

### 3.1. Survey

To investigate the general state of visualization in ERP research, we conducted an online questionnaire using *LimeSurvey Community Edition* v5.4.7 hosted by the *University of Stuttgart*. The survey was open from January to July 2023, and a convenience sample was collected by promoting it through various channels: mailing lists (*EEGLAB, FieldTrip, MNE* communities), social networks *(Twitter, Mastodon, VK, Telegram*, and *Facebook)*, and by speaking at colloquia and conferences. To encourage participation, we raffled 3 *Muse-2* devices (∼300€ each) among those who agreed to participate. We received a total of 425 responses, of which 213 were completed and subsequently analyzed. Since we collected a convenience sample in the first place, we took no further measures to determine why respondents dropped out of the survey.

The survey was approved by the *Ethics Committee of the University of Stuttgart* before it was conducted. Respondents were asked for informed consent as well as to agree to a privacy statement before participating in the study. Additionally, they were informed that they could stop the survey at any time. The anonymous raw data are available in the DaRUS repository (Mikheev, Skukies, et al., 2023a).

### 3.2. Survey structure

The survey was divided into four sections covering different aspects of the respondents’ experiences and opinions. The first page collected basic information about the participants and their experience with plotting. The second page contained questions about the tools they used for analysis, their preferred features in those tools, and their ability to identify different types of ERP plots. The third section asked participants about their familiarity with color mapping and color bar awareness. The final section allowed participants to provide feedback.

Within and across sections, we used a conditional decision-tree structure: certain responses opened additional questions. For example, in the second section, we asked respondents to identify which of the visually depicted ERP plot types they are familiar with. Based on their responses, they then received further questions about the plots they recognized. As a result, some questions received fewer responses than others. For instance, to answer the question “*What did the error bars in your figure represent?*” (85 responses out of 213), respondents first had to positively answer two questions: did they recognize the ERP plot and have they ever used it in one of their papers.

On the survey landing page, we informed respondents of the purpose of the survey, the opportunity to enter the raffle, and the opportunity to opt out of the study. On the second-to-last page of the survey, participants were asked to provide any additional comments or suggestions for improving ERP plotting tools. The final page of the survey contained a link to a separate survey for those who wanted to enter a raffle and/or receive news about this study, a request to share the survey with colleagues, and a short list of publications on perceptual controversies with color maps.

### 3.3. Effect of proficiency

We wanted to understand whether expertise of respondents affects their responses of tool-feature preference. To do so, we first identified a latent *research proficiency factor*, out of our five proficiency-related items: number of papers published, years of experience, current research occupation, self-assessed level of proficiency, and code contribution for any EEG analyses. We used FAMD (Factor Analysis for Mixed Data), a mixture of PCA and MCA methods for continuous and categorical variables, from *R* package *FactoMineR v2.8* (Lê et al., 2008). Using this latent factor, we conducted several univariate linear and logistic regressions with proficiency as an independent variable and with various visualization customs and preferences as dependent variables.

### 3.4. Analysis of free-form naming

We presented respondents with eight ERP plots and asked them to select the ones they recognize. Conditional on this, they were asked to name the recognized plot types in a free form later in the survey.

To analyze the free-form name responses, we first converted them to lowercase, removed unimportant words such as articles and certain terms (“wise”, “like”, “s”, “plot(s)”), and manually eliminated nonsensical or missing responses. Due to the high variability of the responses, we categorized each response into summary names, with summary names identified by manual assessment. For example, when analyzing the ERP plot, we grouped responses containing “erp” and “average(d)” into “averaged erp,” while those containing “erp” and “time” or “time(series)” were classified as “erp timecourse / timeseries”. A full list for each plot type can be found in the provided analysis code (Mikheev, Skukies, et al., 2023b).

### 3.5. Challenges in figure creation

Based on the condition above we asked participants to share the challenges they encountered in creating the respective figures and to provide suggestions on how to improve the tool used to create them. This feedback was collected and made available in full to all tool maintainers if they received more than 5 suggestions.

Figures were generated using *R v4.2.1* with the library *ggplot2 v3.4.1* (Wickham et al., 2016) and *Julia v1.9.1* with the packages *Makie v0.19.8* (Danisch & Krumbiegel, 2021), *UnfoldMakie v0.4.1* (Mikheev, Döring, et al., 2023), *UnfoldSim* v0.1.7 (Schepers et al., 2023). All analysis figures are freely available under MIT license (Mikheev, Skukies, et al., 2023b).

### 3.6. Citations

To count the citations per year for the most popular tools, we used the Web of Science portal. For each software, we selected the most cited paper on the portal and downloaded the data using the following click path: Results cited -> Analyze results -> choose Publication years, Bar chart, Results count by date -> Data rows displayed in table (*Web of Science Core Collection*, 2023). The data was retrieved on 02.11.2023.

## 4. Results

In an online survey, we asked EEG practitioners (n=213) about their opinions and use of ERP visualization tools. The average respondent is currently working in Europe or the USA (Figure 4) on their doctoral degree (37%) in a fundamental field (86%) using scalp EEG (70%). The sample further consists of a mix of postdoctoral researchers (29%), and a considerable number of professors (20%) and industry experts (6%). The median respondent has 6 years of experience with EEG analysis and 3 publications on the topic (Figure 5), while the majority identify themselves as researchers with intermediate experience. More than half of the respondents have published code/software for EEG, MEG, or iEEG analysis methods or contributed to code maintained by others.

**Figure 4.**
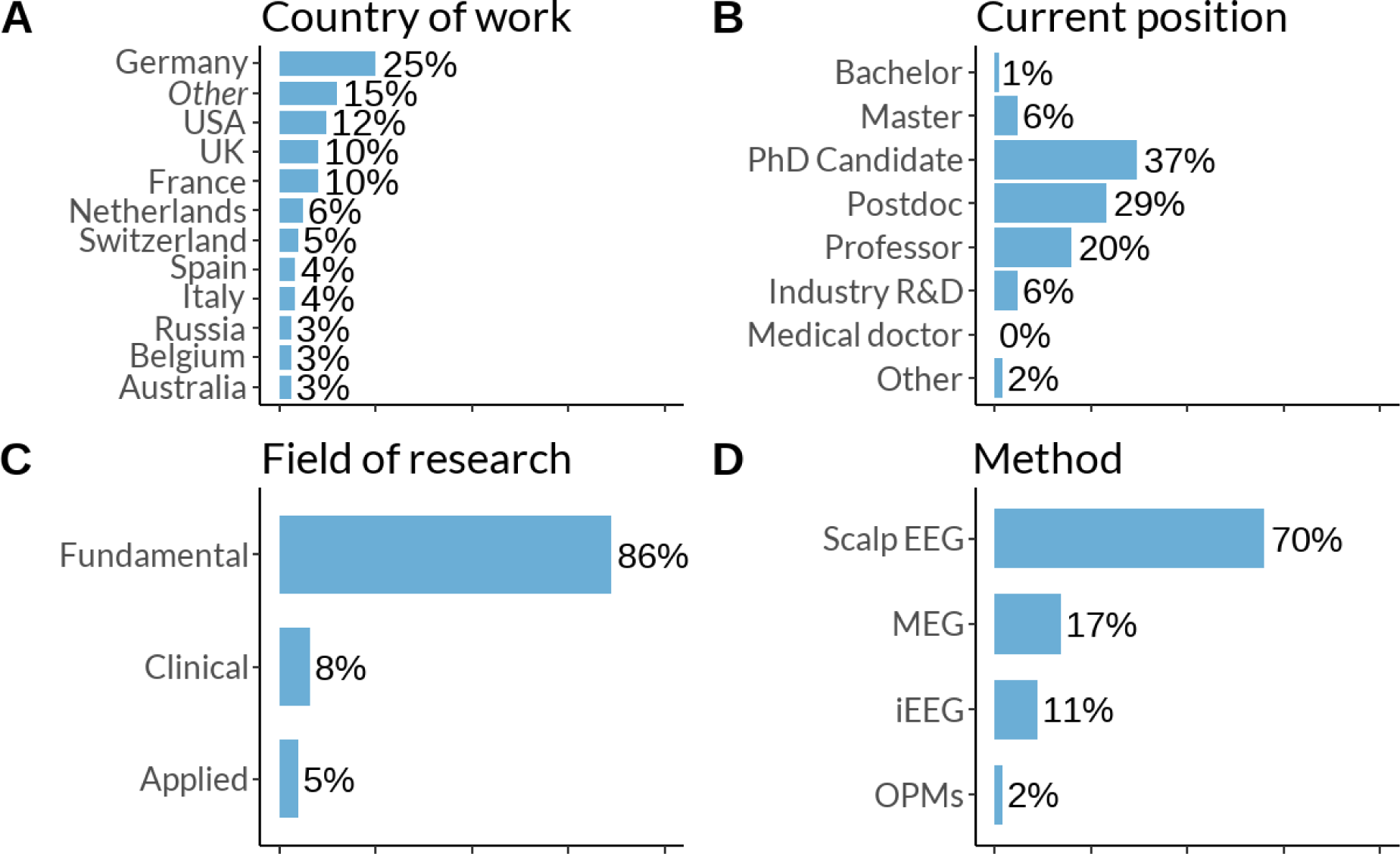
Sample description. D was a multiple-choice question. (N of respondents: A, B, D - 213; C - 212)

**Figure 5.**
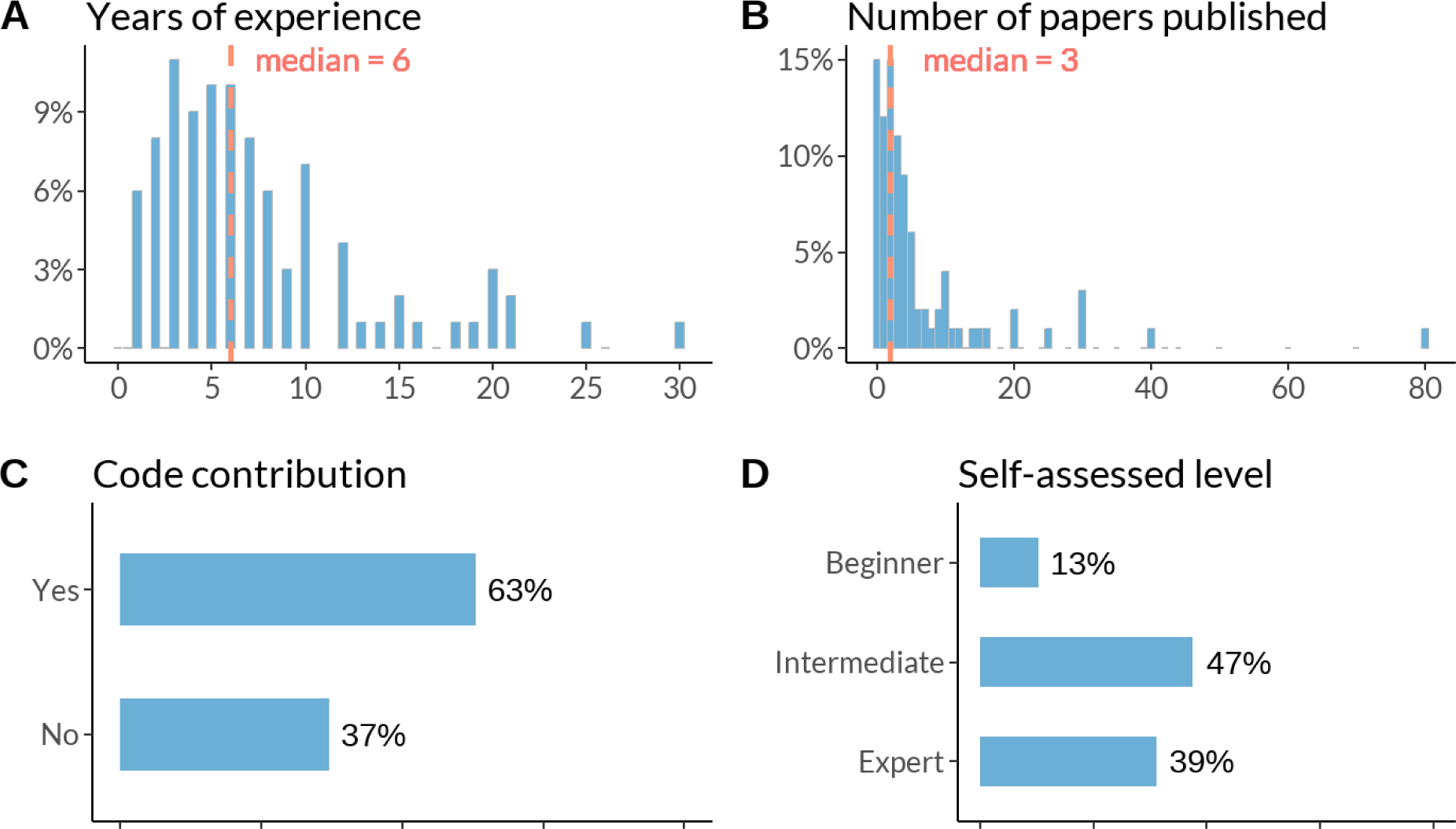
Research experience of respondents (N of respondents: A - 210; B, D, C - 213)

### 4.1. Is there a general naming convention of ERP plot types?

We were interested in how our respondents would name each of the eight ERP visualization types. First, we presented them with depictions of eight plots and asked them to select the ones they recognized. Later in the survey, we additionally asked participants to name these plots in a free form. The attrition of respondents can be seen in Figure 12. The majority of participants have plotted ERP plots and topoplots, while only a tiny portion are familiar with the parallel plot. We further see that e.g. many researchers recognized the channel image and provided a name, but less than half of those have actually plotted it.

Second, we look at the actual suggested names. To do this, we compiled the raw names into aggregated plot names (see Methods). The most popular aggregated plot names, 3 for each plot type, are listed in Table 3. None of the aggregated plot names reached more than 50% of the respective sample. At first sight, this shows that there is little consistency in the naming of ERP plots between researchers. However, at a second look, some names are more closely clustered than others (e.g. 79% majority for the names ‘topoplot’ or ‘topography plot’). A summary and distillation of these results can be found in the Discussion.

### 4.2. Which ERP visualization tools are known and in use?

Most respondents to our survey have experience with MATLAB-based tools (*EEGLAB* - 63%*, FieldTrip* - 46%, and *ERPLAB* - 22%, *Brainstorm* - 16%). The number of users with experience in at least one MATLAB-based tool in our sample is higher than 83%. Custom scripts written in programming languages such as R, Python, Julia, or other languages are commonly used as well (42%). They are followed by the Python-based *MNE* (41%), and a commercial product, *Brain Vision Analyzer* (22%) (Figure 6).

**Figure 6.**
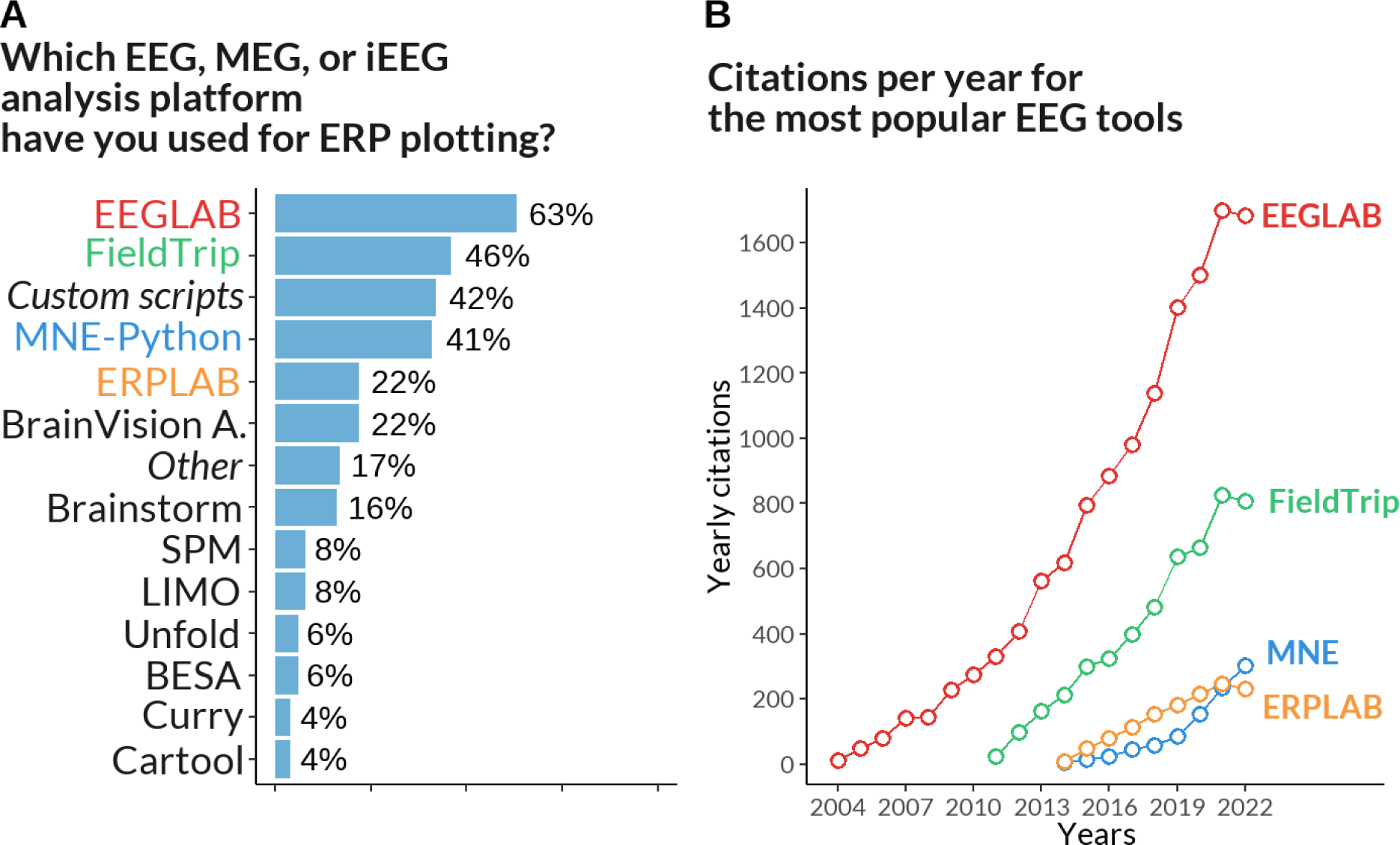
A: User experience with EEG analysis tools (multiple choice, N of respondents - 213). B: The yearly (not cumulative) number of citations for the four most popular tools.

The main caveat to these numbers is that we generally asked about experience with tools, not about the currently preferred tool. As such, participants were able to select multiple answers: one respected researcher outperformed all others by reporting experience with 11 tools.

To complement these results, we obtained annual citations for the four publications of the most popular tools (Delorme & Makeig, 2004; Gramfort et al., 2013; Lopez-Calderon & Luck, 2014; Oostenveld et al., 2011, as indicated “to-be-cited” on their respective toolbox websites). Of this list, EEGLAB was the earliest available tool to analyze EEG data (Figure 7), and is currently the most popular, with yearly citations of over 1600. FieldTrip received two times fewer yearly citations by 2022, while MNE and ERPLAB received four times fewer.

**Figure 7.**
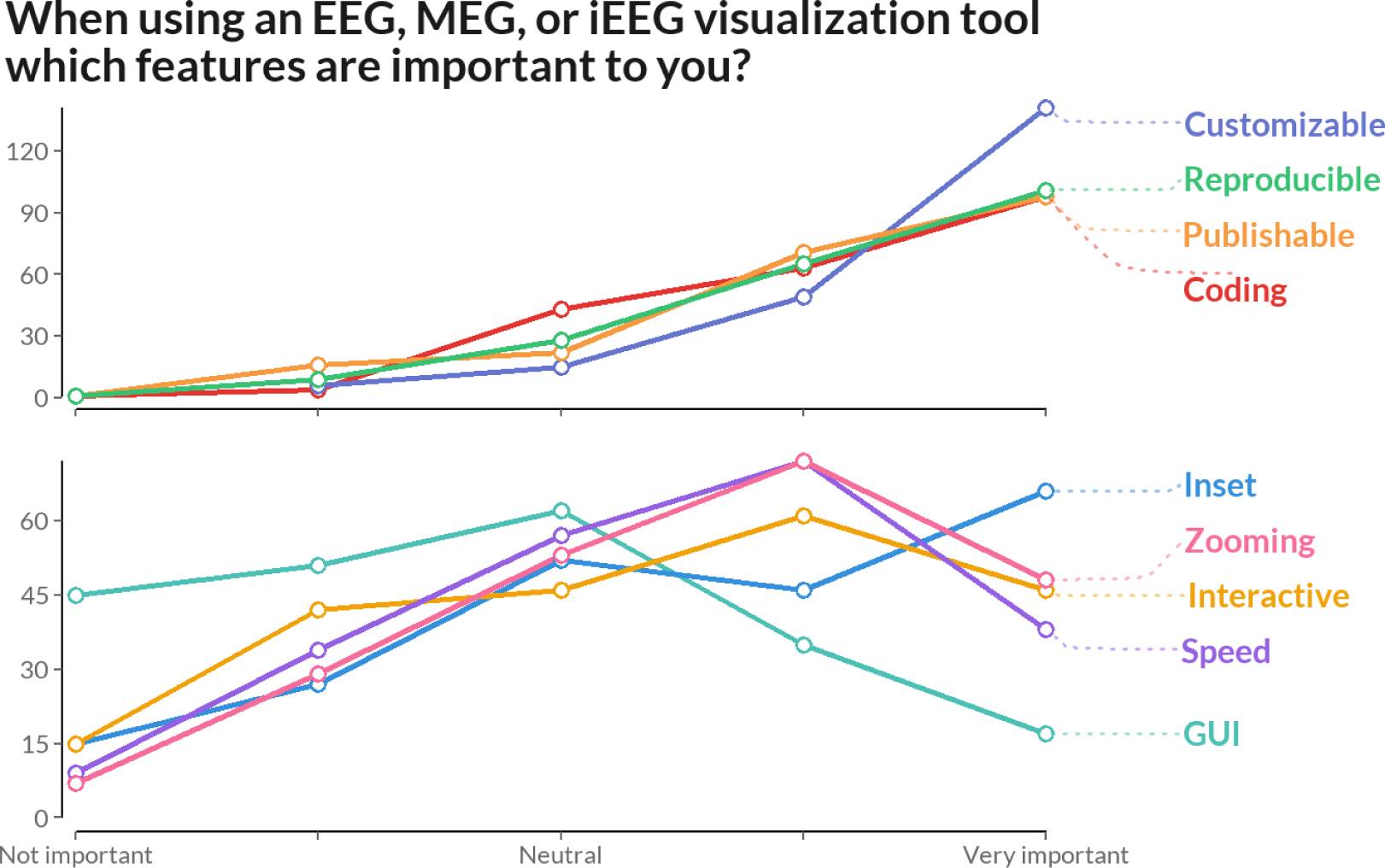
User preferences regarding the features of the EEG analysis tool (N of respondents - 213). To avoid cluttering we split the figure into two panels.

### 4.3. What features are important in current ERP visualization tools and what are potential improvements for tools?

We asked users to rate the importance of various features for EEG visualization tools. Figure 6 shows Likert scale scores ranging from −2 (not important) to +2 (very important) averaged over participants. The majority of users unanimously emphasized the value of customization (flexibility of tweaking plot attributes like colors, line widths, margins, etc.), reproducibility, publishable plots, and generating plots by coding.

Interestingly, some features received mixed responses. For example, the ability for inset subplotting was considered neutral by some users and important by others. A significant proportion of users did not see the importance of speed of plotting and interactive data selection (e.g. direct selection of electrodes in the plot). In addition, EEG practitioners generally did not find it important to generate plots via GUI interfaces.

Additionally, we asked respondents to indicate problems they encountered when plotting a figure using their favorite tools. For example, when it came to creating the ERP plot, 32% of respondents indicated that they struggled with adding uncertainty to their plots. For ERP and channel images, sorting was the most common problem. Three the most mentioned problems for each plot are summarized in Table 2. We omitted parallel plots due to lack of responses.

**Table 2.**
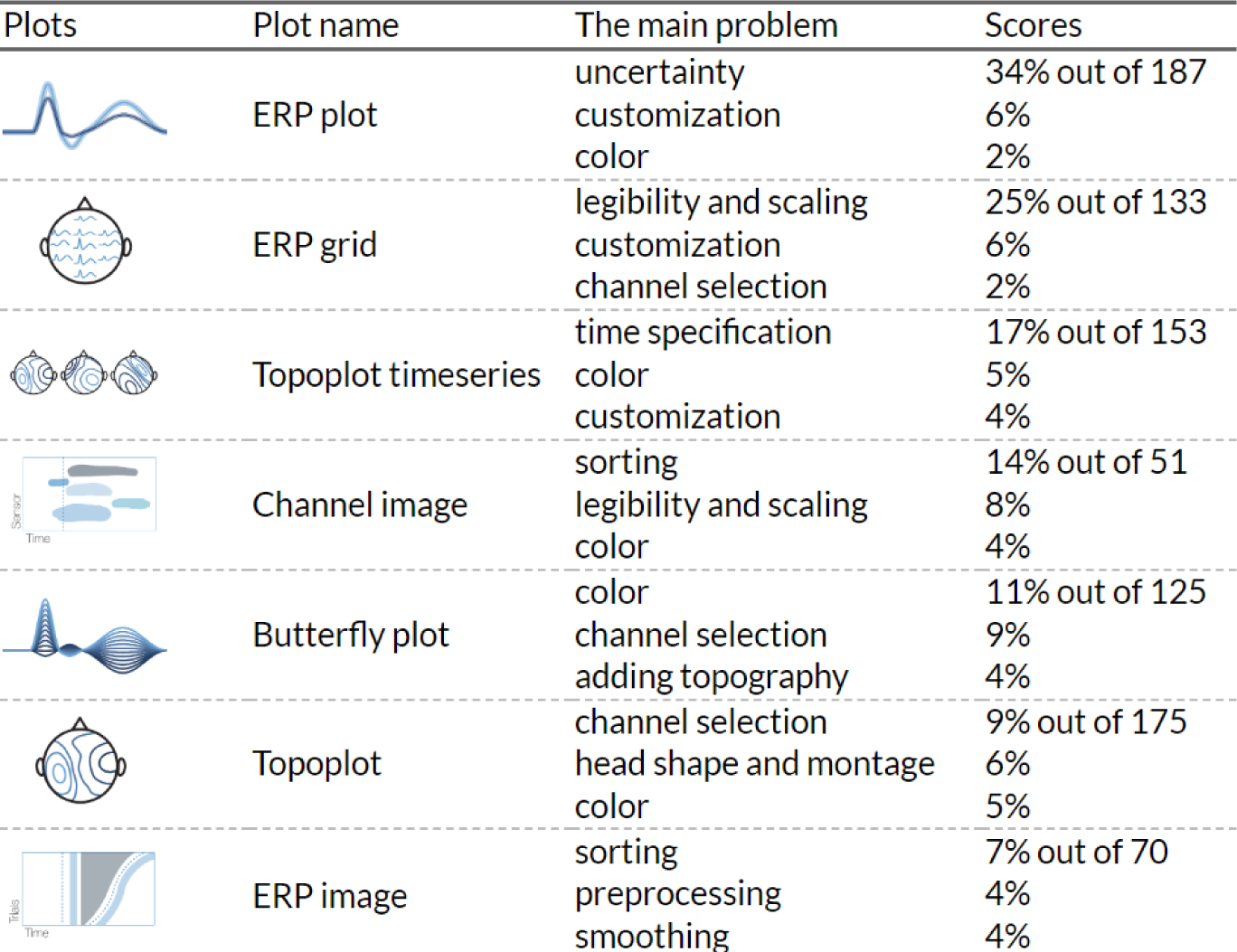
The most common plotting problems for each ERP visualization type.

| Plots | Plot name | The main problem | Scores |
| --- | --- | --- | --- |
|  | ERP plot | uncertainty<br>customization<br>color | 34% out of 187<br>6%<br>2% |
|  | ERP grid | legibility and scaling<br>customization<br>channel selection | 25% out of 133<br>6%<br>2% |
|  | Topoplot timeseries | time specification<br>color<br>customization | 17% out of 153<br>5%<br>4% |
|  | Channel image | sorting<br>legibility and scaling<br>color | 14% out of 51<br>8%<br>4% |
|  | Butterfly plot | color<br>channel selection<br>adding topography | 11% out of 125<br>9%<br>4% |
|  | Topoplot | channel selection<br>head shape and montage<br>color | 9% out of 175<br>6%<br>5% |
|  | ERP image | sorting<br>preprocessing<br>smoothing | 7% out of 70<br>4%<br>4% |

**Table 3.**
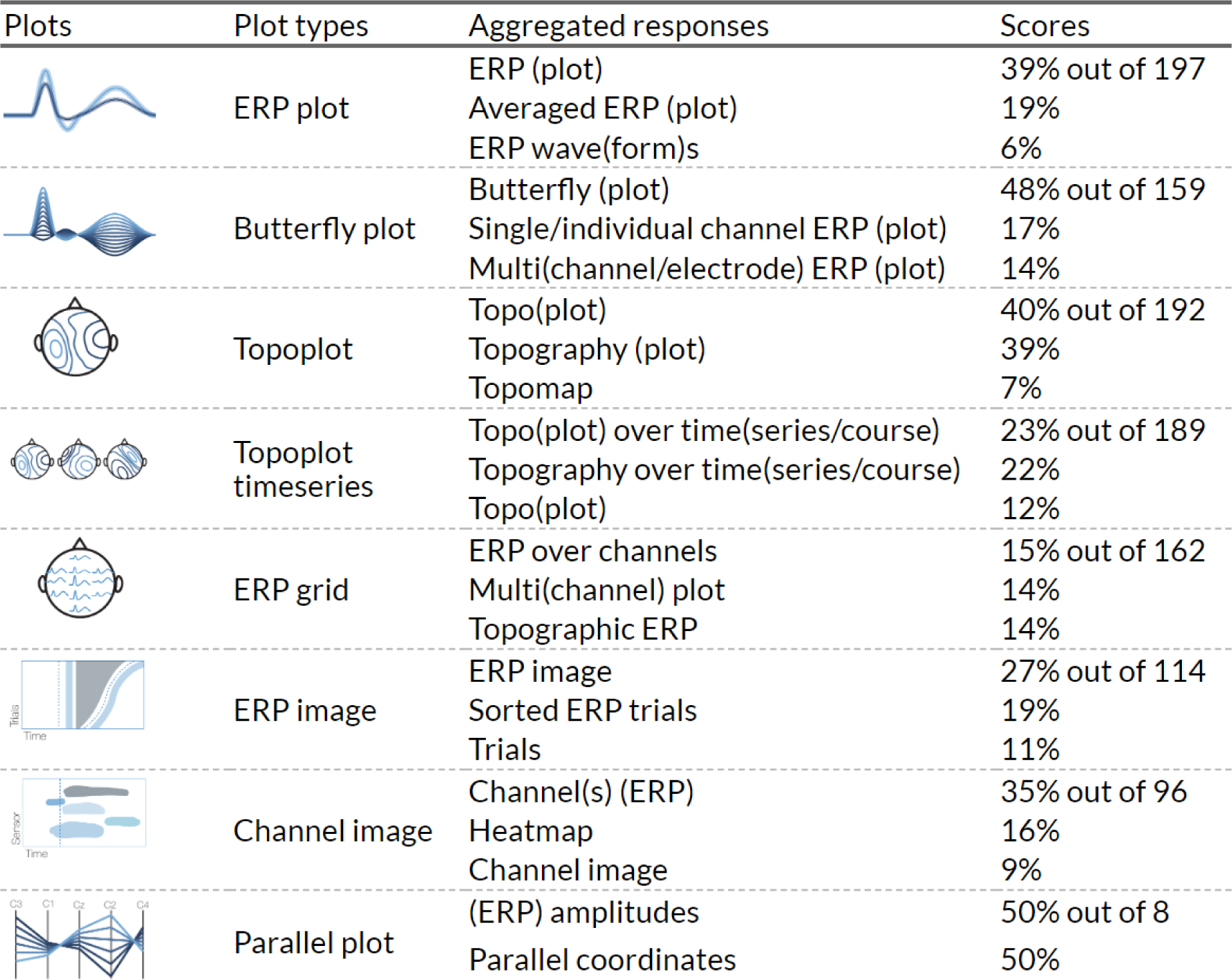
The most popular plot names for eight types of ERP visualization. The names were aggregated. To illustrate the notation: e.g. “ERP (plot)” means that response like “ERP” and “ERP plot” were united, “time(series/course)” means that instances “time”, “timeseries”, “time series”, “timecourse” and “time course” were combined. The word “over” in some names is also an aggregation including all other omitted prepositions like “by” or “with”.

| Plots | Plot types | Aggregated responses | Scores |
| --- | --- | --- | --- |
|  | ERP plot | ERP (plot) | 39% out of 197 |
|  |  | Averaged ERP (plot) | 19% |
|  |  | ERP wave(form)s | 6% |
|  | Butterfly plot | Butterfly (plot) | 48% out of 159 |
|  |  | Single/individual channel ERP (plot) | 17% |
|  |  | Multi(channel/electrode) ERP (plot) | 14% |
|  | Topoplot | Topo(plot) | 40% out of 192 |
|  |  | Topography (plot) | 39% |
|  |  | Topomap | 7% |
|  | Topoplot timeseries | Topo(plot) over time(series/course) | 23% out of 189 |
|  |  | Topography over time(series/course) | 22% |
|  |  | Topo(plot) | 12% |
|  | ERP grid | ERP over channels | 15% out of 162 |
|  |  | Multi(channel) plot | 14% |
|  |  | Topographic ERP | 14% |
|  | ERP image | ERP image | 27% out of 114 |
|  |  | Sorted ERP trials | 19% |
|  |  | Trials | 11% |
|  | Channel image | Channel(s) (ERP) | 35% out of 96 |
|  |  | Heatmap | 16% |
|  |  | Channel image | 9% |
|  | Parallel plot | (ERP) amplitudes | 50% out of 8 |
|  |  | Parallel coordinates | 50% |

### 4.4. What does the EEG community think about some of the controversies surrounding ERP visualization?

Previously, we identified four visualization controversies and asked participants to provide some input on specific aspects: baseline period visualization, proportion of electrodes analyzed, polarity notation and interpretation of topoplot timeseries.

First, participants most commonly answered to show a baseline period of 200 ms (45%) or 100 ms (22%) prior to event onset (Figure 7A). Further 48 (∼30%) of the respondents rather provided additional explanations: 25 said that it depends on the experimental design, 11 noted that it should be the same as the baseline correction, 9 said that it depends on the trial duration.

Second, as shown in Figure 7B, most respondents (38%) used all recorded electrodes for analysis (c.f. Figure 2 with marginal histograms in Supplementary material).

Thirdly, a vast majority (82% of 199 participants) preferred to plot positive voltage upward on the Cartesian graph (Figure 8). Comparing different research fields, we found that researchers working with language and in cognitive control and attention seem to be more likely to choose negative up. However, even in those fields, the majority preferred positive up.

**Figure 8.**
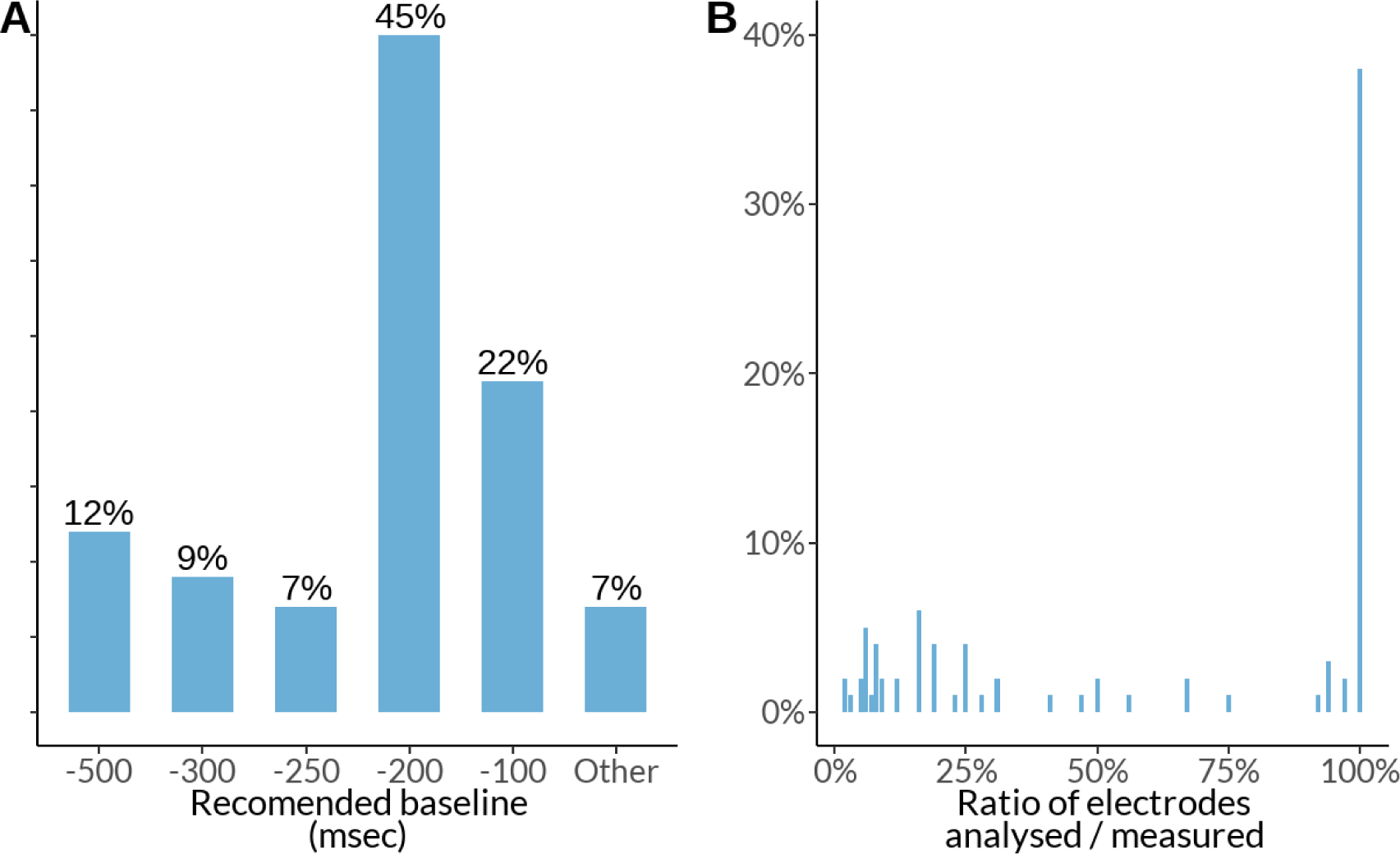
A: Recommendations for baseline period visualization in ERP analysis (N of respondents - 163). B: Ratio of analyzed vs. recorded electrodes (N of respondents - 208).

**Figure 9.**
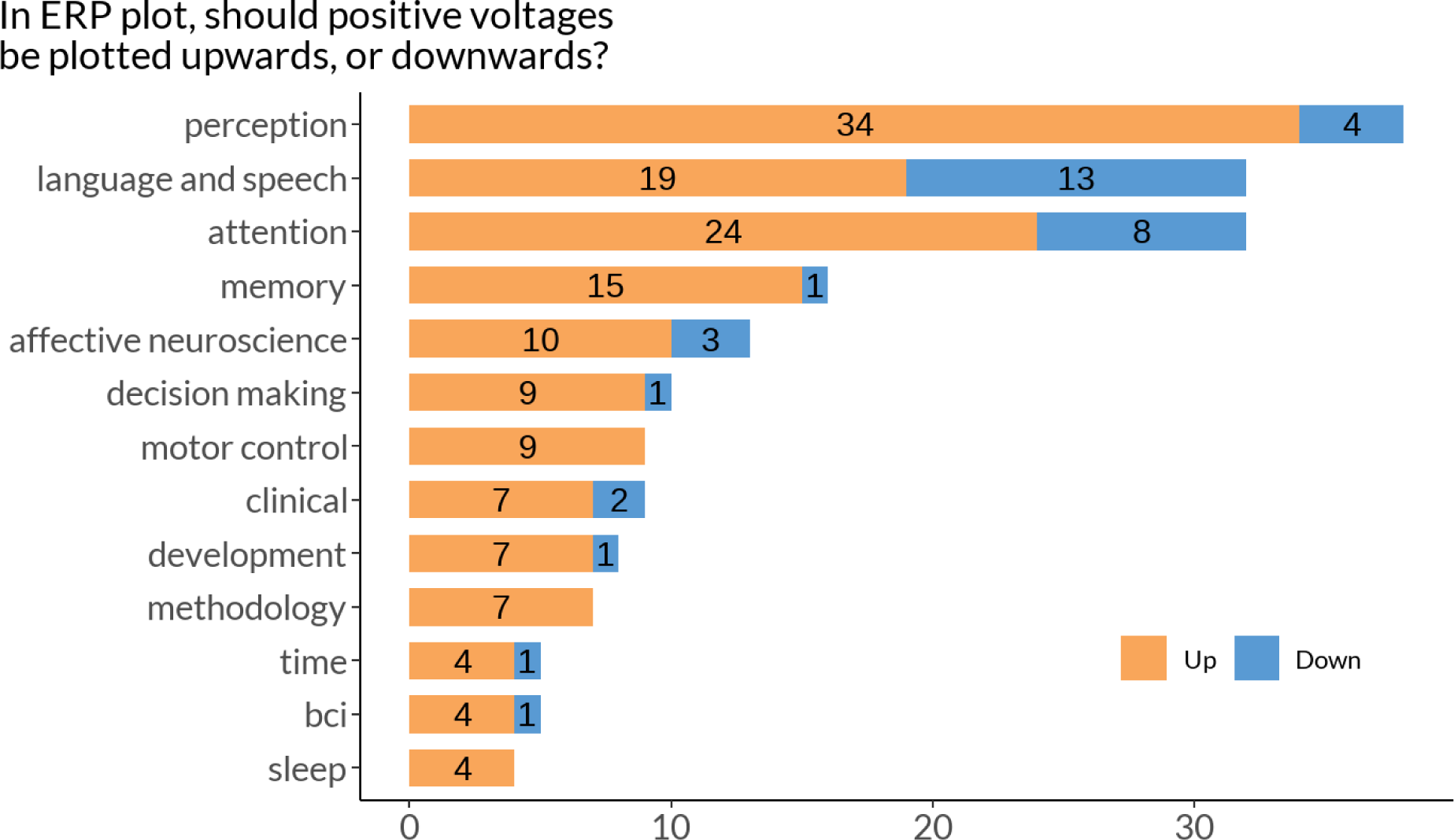
User preferences concerning voltage orientation by field of research (N of respondents - 188)

**Figure 10.**
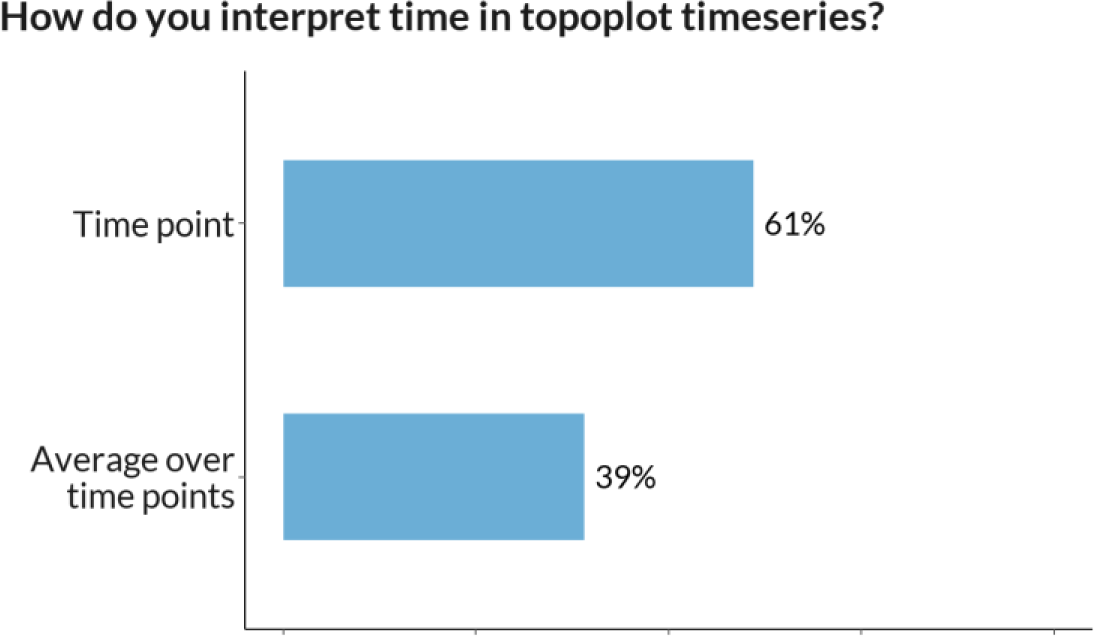
Interpretation of time in topoplot timeseries (N of respondents - 185)

Lastly, in a topoplot series, we found that 61% of the 185 respondents understand each topoplot as a snapshot of scalp electrical activity at a particular time point. While 39% interpret it as an average over a time interval. Thus, it is a potential source of confusion and to avoid this, researchers should clearly label time interpretation on their topoplot series.

### 4.5. How well are ERP researchers aware of some of the perceptual issues in visualization?

To test visualization literacy, we asked respondents about their awareness of issues related to color maps and error bars.

Out of 213 respondents, 39% were unaware of the perceptual controversies regarding color maps. This shows that more emphasis needs to be placed on such issues in educational curricula, and perhaps software developers should encourage users to use better color maps by setting appropriate defaults.

Next, out of all respondents who had previously published an ERP plot (n=157) 40% did not include error bars (Figure 11A). This percentage is worse than in previous studies for non-timeseries plots (∼20%) (Allen et al., 2012). Note that Allen et al. did not consider timeseries, where uncertainty visualization is more challenging, and did not conduct a survey; they counted visualizations in papers.

**Figure 11.**
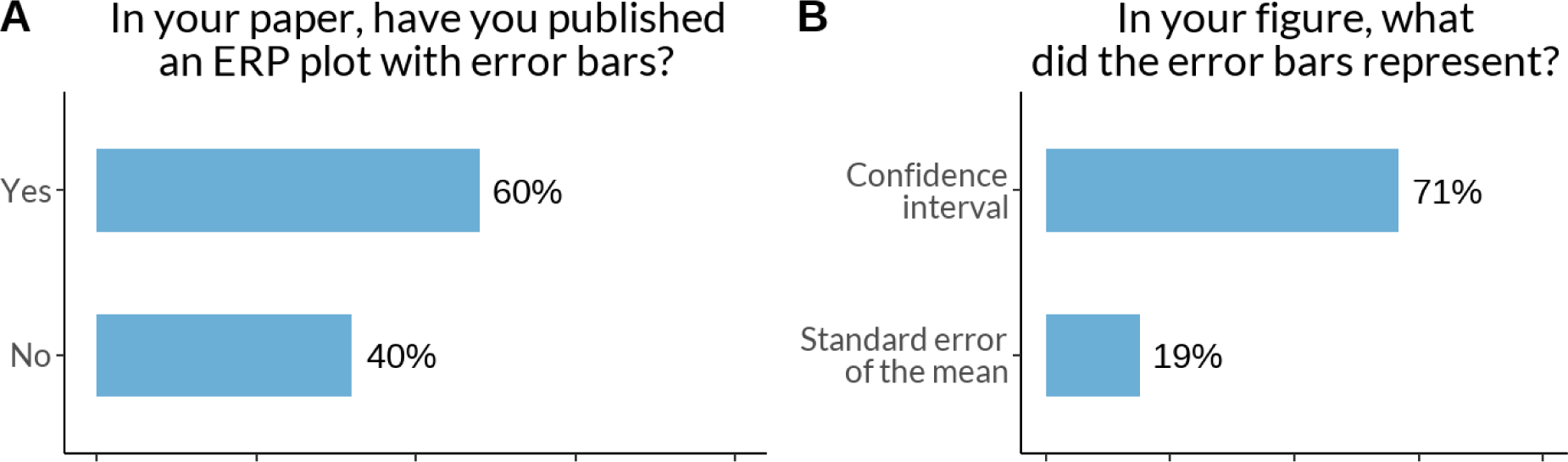
The error ribbons on a line graph and how researchers interpret them (N of respondents: A - 157, B - 85)

**Figure 12.**
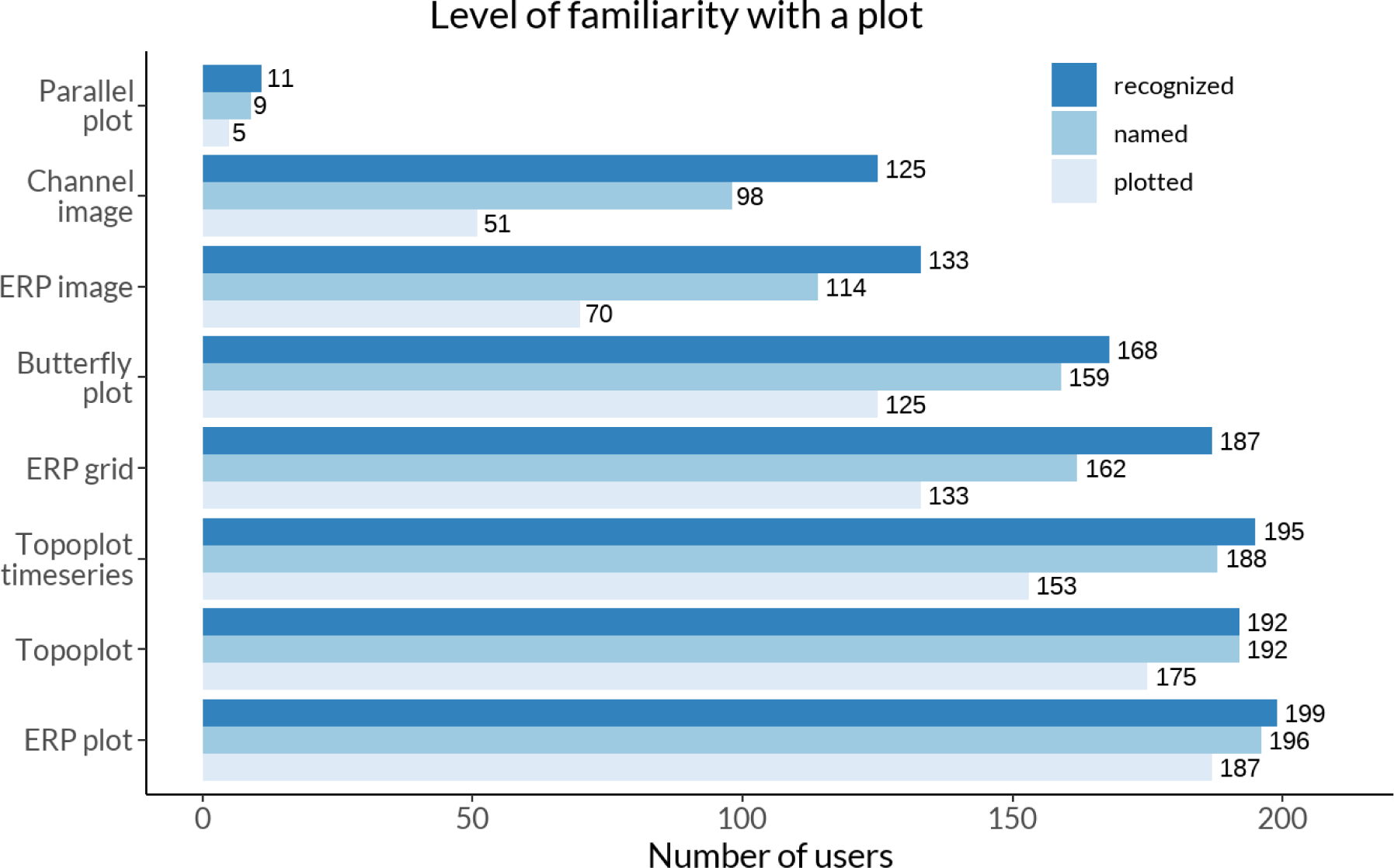
Number of respondents who recognized, named, or had previously plotted the respective plot types. (N of respondents - 213)

We further asked for the depiction of uncertainty in heatmaps, which is notoriously difficult to do. One solution is to use 2D color maps (aka bivariate color maps or cross-maps). Bivariate color maps were first used by the US Census Bureau in 1974 to describe the age of older Americans and the population size of counties on the US map (Meyer et al., 1975). Later Correll picked up on that to graphically show the relation between statistical power and raw effect size (Correll et al., 2018). We were interested in whether researchers are aware of this option: 68% of the 213 respondents indicating that they had heard of it. Interestingly, more than 76% (of those being aware) would like to use 2D color maps for uncertainty visualization in the future if they were easily available in their software packages. However, to our knowledge 2D color maps are only available in a few selected analysis/visualization packages (e.g., *EEGVIS* or *FieldTrip* with custom code).

### 4.6. Does proficiency in EEG correlate with opinions towards ERP visualizations requirements?

In this section, we wanted to test how proficiency affects preferences, habits, and awareness of ERP visualization.

First, we explored if there was a correlation between proficiency and preference for a specific research tool. Out of 27 tools, 10 tools plus custom scripts as a unique category were selected for analysis based on the criteria that more than 10 people used this tool. 10 logistic regressions were run with 4 significant estimates, indicating a potential correlation between higher proficiency of researchers and the experience with using any such tool: *EEGLAB (β=0.265, p=0.006)*, *ERPLAB (β=0.20, p=0.042)*, *FieldTrip (β=0.19, p=0.023)*, and *LIMO (β=0.34, p=0.021)*. Note that experience typically correlates directly with age, so the correlations found could simply be an effect of how old a given toolbox is. Please find all results in Supplementary Table 1.

Second, three logistic regressions were conducted to assess attitudes toward polarity notation, awareness of color map controversies, and willingness to use bidimensional color maps. We found only one significant correlation: with higher proficiency, researchers were more likely to know about the perceptual controversies surrounding color map controversies *(β=0.34, SE=0.1, p<0.001)*.

Third, using Spearman rank correlations, we observed that there is a positive significant correlation between proficiency and the number of tools used *(ρ=0.31, p<0.001)*, and the ratio of channels analyzed to channels recorded *(ρ=0.26, p<0.001)*. While the first result is rather obvious, the latter is less so: more experienced researchers tend to keep all recorded channels in the analysis.

Finally, the same Spearman rank correlations were used to analyze associations of proficiency with indication of importance of software features. Out of eight software features, only the importance of “generating a figure by coding” correlated significantly with proficiency, showing a small positive relation *(β=0.17, ρ=0.013)*. See more in Supplementary Table 2.

Lastly, we want to stress that none of these findings show any causal effects, they all depend on a biased sample and the effect sizes are quite small. We also did not correct for multiple testing, as taking such p-values at face value is a fool’s errand in the first place.

## 5. Discussion

In this study, we explored the current state of ERP visualization practice in the EEG community and tried to understand the challenges faced by practitioners. Our research questions focused on identifying popular ERP visualization tools, determining which features of these tools are important for practitioners, investigating controversies in ERP visualization, and assessing the awareness of researchers regarding visualization issues.

### 5.1. Recommendations on the plot naming

No consensus plot name was found for most ERP plots. The strongest results were for the “topoplot”, which combined with the “topographical plot” received more than 80% of the responses. For some, even after aggressive aggregation, no common plot name remained, as in the case of “ERP grid plot”.

Even though it became clear that no consistent naming scheme exists, there are still several reasons why it would be very helpful to define one. 1) Discoverability of functions. A common lexicon is useful for naming visualization functions. The research software should be intuitive and accessible to new users. Function names should therefore be easy to find in the documentation. 2). Consistent reporting. It is helpful to have clear options in the visualization section when standardizing reports such as ARTEM-IS, which is beneficial for reproducibility and meta-analysis. 3) Education. It is easier if teaching materials and resources do not use different names for the same visualization.

For each plot type name, we carefully weighed the popularity of the proposed names among researchers, uniqueness compared to other commonly used plot types, and their logic and memorability.

- ERP plot. With 40% of the votes this name was the most popular among EEG practitioners. The runner up suggestions, such as “averaged ERP” seemed a bit redundant (the great majority of ERPs are averaged) or did not receive many votes (e.g. “ERP waveform plot”). While a typical “ERP plot” is technically a type of line plot, it is not referred to by this name among EEG researchers (3%).
- Butterfly plot. The name “butterfly” (48%) doesn’t describe the underlying plot type or data, such as names “multi-channel line plot” or “multi-channel ERP plot”. However, it is still a catchy and popular description of the resulting pattern as the mirror symmetry of the lines is reminiscent of the symmetry of a butterfly. One question remains: what is the difference between a butterfly plot and a multi-line ERP plot? We suggest that a butterfly plot is one instance of a multi-line ERP plot, which refers to the plotting of multiple <u>channels</u>, whereas the plotting of multiple single <u>trials</u> or multiple <u>subjects</u> could be referred to with a qualifier, e.g. “single-trial/multi-trial ERP plot” or “multi-subject ERP plot” whenever appropriate.
- Topography plot (short “topoplot”). The topography plot is a 2D projection and interpolation of the 3D distributed sensor activity. The name stems from physical geography, but instead of height, the contour lines represent voltage levels. Respondents were fairly unanimous that either topography plot (40%) or topoplot (39%) should be chosen. We decided to promote both names, as topoplot is a convenient and unequivocal nickname to topography plot.
- Topography timeseries (short “topoplot series”). Technically, a multiple miniature topography plot, most of the suggested names included prepositions (e.g. “by”, “on”, “at”), but we decided to leave them out for the sake of brevity. The most popular result was “topoplot/topography timeseries” (23%/22%). The runner up “topoplots” is ambiguous with the topography plot introduced above, so we decided to add ‘timeseries’ for clarity. This retains the link to traditional topography plots but highlights the differences.
- ERP grid (plot) This is the most challenging plot to name for EEG practitioners, as the community could not agree on a unique, descriptive plot title. We chose “ERP grid” because the multiple miniatures of the ERP plot in this figure are typically arranged according to electrode positions, mimicking a grid across the scalp. Alternative names were “ERP montage / scalp”, “ERP channels” or “multi-channel ERP”, “multi-axis ERP” or “topographic ERP / array”. However, the terms were either not well known (such as “montage”, “array”), misleading (“topographic” - there are no height lines, “multi-axis” - a name that typically refers to e.g. two y-axes in one plot), or potentially confusing (“multi-channel ERP” could be confused with either a “butterfly plot”, or an “ERP plot” averaged over multiple channels). We therefore suggest “ERP grid” as the name for this plot type, even though we are aware of the issues with proposing yet another standard (Munroe, 2013)
- ERP image This plot is technically a heatmap, but most commonly known as an “ERP image” (27%). It was proposed under this name by Jung et al (Jung et al., 1998).
- Channel image The two most popular options, “channels ERP” (35%) and “heatmap” (16%), are both misleading: “channel(s) ERP” could be confused with the “butterfly plot”, and “heatmap” is too general and can be confused with the “ERP image” plot type. For consistency and simplicity, we therefore used the third most popular choice (9%): “channel image”.
- Parallel coordinate plot (parallel plot). This plot type is very rarely used in the EEG literature (one example in Caat et al., 2007), which also reflects in the answers to our survey: only 8 respondents recognized this plot type. We therefore decided to keep the formal name from the visualization literature, coined in 1885 (Moustafa, 2011).

### 5.2. Popularity of ERP visualization tools

This distribution of software experience is useful for inferring trends in software experience. Users should know what tool to learn and software developers where to invest time and effort. Furthermore, the selection of software toolboxes or versions may contribute to variations in the results (Kabbara et al, 2023 on 3 MATLAB-based tools), thus knowing the most popular tools helps to be aware about possible methodological biases in the field.

According to our survey, most EEG practitioners (82%) have experience with MATLAB-based tools, including *EEGLAB, FieldTrip, ERPLAB* and *Brainstorm*. The Python-based *MNE* and commercial software *Brain Vision Analyzer* also showed significant usage, as well as custom scripts written in programming languages such as *R, Python, Julia.* However, experience with a tool does not directly reflect in citing the tool. About 1.5 times more respondents had experience with EEGLAB than with its closest competitor, but the yearly citations show a gap of 2 to 4 times. This is either an indication that our question asked for experience and not application, or a sampling bias in our survey.

It is interesting to compare these results with the survey from (Hanke & Halchenko, 2011)^4^ on research software in neuroscience. This allows us to make some careful comparisons of what changed in EEG software popularity during the last 12 years (Table 4). In 2011 the first three leading positions were taken by EEGLAB, Fieldtrip and Brainstorm.

**Table 4.** Comparison of the usage of some softwares in Hanke & Halchenko and our study.

| Tool<br>Paper | EEGLAB | Fieldtrip | Brainstorm | Brain Vision<br>Analyzer | MNE | ERPLAB |
| --- | --- | --- | --- | --- | --- | --- |
| 2011 (Hanke & Halchenko)<br>N= 181 | 92% | 44% | 31% | 31% | 17% | 1 user |
| 2023 (ours)<br>N=213 | 63% | 46% | 41% | 22% | 41% | 22% |

Brainvision Analyzer was fourth, MNE only seventh, while ERPLAB got only one user. We can observe the following trends: 1) Users still have the most experience with MATLAB-based toolboxes; 2) Brainstorms popularity decreased and ERPLAB took its place; 3) MNE moved from seventh to third place (excluding custom scripts). Note that the survey sampled all of neuroscience (we used numbers only for M/EEG researchers) and asked for what they use in their research activities.

Recently, the EEGManyPipelines project (Cesnaite, 2023) asked 168 teams to test eight hypotheses based on the same provided data. They preliminarily released the software packages the teams used. Interestingly, MNE was the most popular tool (44 teams), only then followed by EEGLAB (42) and Fieldtrip (31). Note that this sample may be biased toward more experienced and methodologically oriented teams.

Together with the citation pattern (Figure 7), this leaves us with four resources for assessing current tool experience and usage. It seems that MNE-Python is on the rise, with currently the fastest growth rate, and already has the most users among the *EEGManyPipeline* teams. However, by far the largest share of experience and usage is combined in MATLAB-based tools, especially EEGLAB.

### 5.3. Important tool features

We found that practitioners often faced specific problems when generating different ERP visualizations. For instance, adding uncertainty to ERP plots, styling and color in butterfly plots, channel highlighting in topoplots, and legibility and scaling in ERP grids were among the most common challenges. These results partially mimic other literature: representing uncertainty was discussed as one of the most challenging research problems in scientific visualization already in 2004 (Johnson, 2004).

Moreover, our survey indicated that customizability, reproducibility, generating plots by coding and publication-ready plots were highly valued features for ERP visualization tools. However, there was skepticism regarding interactive data selection methods. EEG practitioners, and experts even more strongly, also expressed a preference for generating plots through scripting rather than using GUI interfaces. These insights can provide guidance for improving existing visualization tools and developing new ones that align with the preferences and requirements of users.

### 5.4. Community opinions on ERP visualization controversies

Next, we investigated four controversies surrounding ERP visualization.

1. Our results showed that electrode selection varied across researchers, with only about 38% using all recorded electrodes for analysis. Researchers who subselect channels should be aware of potential issues with expanding researchers’ degrees of freedom, circular analyses (Lyon, 2017) or even cherry-picking. If a-prior recommendations from the literature (e.g. Rossion & Jacques, 2008 for the N170) cannot be used, it is also possible to carefully use data-driven methods (Brooks et al., 2017) or, of course, simply use all channels in a mass-univariate analysis (Ehinger & Dimigen, 2019; Fields & Kuperberg, 2020; Groppe et al., 2011; Pernet et al., 2011; Smith & Kutas, 2015).
2. Regarding the visualization of baseline duration, 45% of respondents suggested 200 ms and 22% suggested 100 ms as optimal, followed by 12% for 500 ms. We found baselines as short as 100 ms surprising, as gauging the noise level seems difficult (but no empirical study is available as of now). We are further quite sympathetic to the respondents, who argued that the period should be adapted to the length of the whole trial. Consequently, we recommend showing a baseline of at least 200 ms, and, as some participants pointed out, at least the size of the baseline-correction period you chose.
3. The majority of respondents (82%) preferred positive voltages to be displayed up on the Cartesian coordinate system, contrary to the historical practice to sometimes show negativity upward. Thus, “positivity up” gathers support and can be regarded as the predominant way in current practice.
4. About 60% of respondents perceive each topoplot as a snapshot of scalp electrical activity at a particular time, while the other 39% perceive it as an average of a time interval. We call to attention both authors to label their plots, but also readers to read carefully what is actually depicted. There is no empirical study of the practical differences between these two approaches, but it is reasonable to think that single time snapshots will have weaker signal-to-noise ratios and be more affected by low-pass filtering.

### 5.5. Awareness about visualization problems

For RQ4, we assessed researchers’ awareness of visualization issues specifically related to color maps and uncertainty visualization. A significant proportion of respondents (39%) were not aware of any perceptual controversies related to color maps. In addition, approximately 40% of those who published an ERP line plot did not include error bars, which are important for accurate data interpretation.

Our results are in line with direct studies of the prevalence of jet color maps in EEG. Approximately 40% of our respondents weren’t aware of this issue in 2023 and ∼60% of published (time-frequency) figures used jet color maps in 2020 (Cooper et al., 2021). While generating more awareness is helpful, another idea is to implicitly discourage the use, by defining better defaults. For example, as of 2023, EEGLAB still uses the *jet* color map as the default. Some discussion was started in 2022 (Delorme, 2022) to change the default to *turbo* (improved but still problematic rainbow color map), but so far, no further progress has been made.

In general, the field of EEG shows one of the highest usages of rainbow color maps in the literature (Table 5). Unsurprisingly, most awareness can be found in the field of visualization, where usage of rainbow color bars per year in papers dropped from 39.6% in 1993 to 1.9% in 2020 (Golebiowska & Çöltekin, 2022).

**Table 5.** The usage of rainbow color map in 6 different fields.

| Field | Years and N of papers | Rainbow palette usage | Study |
| --- | --- | --- | --- |
| EEG (time-frequency plots) | 2000-2020 (n=1652) | 74% of figures on average used the rainbow color palette and its derivatives with a peak in 2016 (~85%), and decrease to ~60% in 2020 | (Cooper et al., 2021) |
| Functional Neuroimaging (fMRI and PET) | 1996-2009 (n=3993) | 15.2% of all figures are in the rainbow scale (RBS)<sup>5</sup> | (Christen et al., 2013) |
| Geoscience | 2005, 2010, 2015, 2020 (n = 2638) | 55% of all papers contained at least one visualization using rainbow or red-green color schemes<sup>6</sup> with the peak in 2015 (61%) and decrease in 2020 (55%). | (Westaway, 2022) |
| Fisheries acoustics (hydrology) | 2009-2019 (n=100) | 16% of papers used rainbow color maps.<sup>7</sup> | (Blackwell et al., 2020) |
| IEEE Visualization conference proceedings | 1990-2020 (n=3057) | The number of papers that contain a figure with rainbow colors decreased from 39.6% in 1993 to 1.9% in 2020 | (Golebiowska & Çöltekin, 2022) |
| Remote sensing and planetary science | 2016-2020 (n=155, 150) | 48% and 64% of papers used rainbow color map | (Golebiowska & Coltekin, 2022) |

These findings suggest that to ensure reliable and meaningful representations of data, we need to increase awareness of visualization best practices not only among scientists, but also among software developers.

### 5.6. Correlation with proficiency

We found that researchers with more experience are familiar with more analysis toolboxes and generally had more experience with either EEGLAB, ERPLab, FieldTrip or LIMO compared to other toolboxes. Whether specialization, popularity, or the age of the toolbox plays a decisive role in this is not entirely clear.

Experts further consider the generation of plots by code as a more important feature compared to less experienced researchers. They are also more often aware of the perceptual issues of color bars and tend to keep more recorded electrodes during their analysis.

The limitations mentioned in the method section are worth repeating: the effects were small, are based on a convenience sample and should be interpreted with care.

### 5.7. Best practices for ERP plots visualization for researchers

There are already some recommendations from COBIDAS for ERP plot visualization (Pernet et al., 2020):

- label all axes and report all units. Direct labels are preferred.
- Plot all channels and conditions. As we can see in RQ5 more experienced researchers tend to analyze more recorded electrodes.
- Show error measures (CI or SD) and label them.

However, we would like to expand this list with insights from our study and practice:

- Pay attention to color maps, they should be scientific: perceptually uniform, intuitively ordered, and colorblind friendly (Moreland, 2016).
- Be aware about where to use diverging and sequential color maps. The former are used when there is a meaningful middle value, while the latter are for scales with continually increasing or decreasing values (Muth, 2021a, 2021b).
- Plot positivity upward.
- Clearly label topoplot timeseries with what each topoplot represents. E.g. the range bracket notation [0.12-0.14) would indicate that the samples are taken from 0.12 (including 0.12) to 0.14 (excluding 0.14).
- Channel and ERP images typically use diverging color maps. For circular plots, such as the position-color encoding in a butterfly plot, use circular color maps. (Figure 3B).
- If you use topoplot timeseries, keep heights of color limits and contour lines identical across subplots (Ehinger, 2016).
- Include at least the baseline period which was used for the baseline correction.
- Explore 2D colormaps that can indicate both voltage and uncertainty. You can find detailed guides for 2-dimensional color maps here (Hengl, 2003).

We further recommend reading the excellent paper from Allen et al with suggestions about visualization of axis, uncertainty, annotation, and color (Allen et al., 2012). Also read this paper about the importance of adding uncertainty in a plot instead of showing brute summary statistics (Rousselet et al., 2016).

### 5.8. Suggestions for software developers

While there are already some recommendations for research software development, they are mostly of a rather general nature (Gewaltig & Cannon, 2014). As such, additionally to our main survey focus, we collected raw feedback on toolbox usage (usage complaints, suggested features, and general recommendations on plotting) and sent them to the six respective toolbox developers by mail or via Github issues. Here we suggest accumulated recommendations concerning software development in the field of electroencephalography visualization in general:

- Use intuitive names for functions.
- Make scientific color maps the default option.
- Make upward polarity notation the default option.
- Encourage users to give labels to topoplots which indicate whether a single time-point or an average of time-ranges is used (e.g. using the range bracket notation).
- Make error bars and error ribbons easy to use and nudge users to always include them.
- Use plotting backends that allow for flexible customization by the users.
- Ensure the plots can be made reproducible, especially if the users can edit them via a GUI.

### 5.9. Limitations

This study has a number of limitations that reduce the generalizability of our findings. First, we surely have a sample bias, meaning that the people who participated do not fully represent the entire population of scientists using EEG. For instance, the majority of our sample is made up of EEG researchers working in rich, western countries with a long and well-established tradition of scientific research: 76% of our participants are from Europe (with 49% are from Western Europe) and 13% are from North America. Scientists working in Africa, Asia, and South America are not well represented in our study (1-3% each). Further, 86% of our respondents are from fundamental research, shadowing researchers from the medical or applied fields. The survey was conducted in English, while in other languages the results, especially concerning plot naming, could be different. All these reasons may affect the extent to which we can generalize our findings to all EEG practitioners.

In addition, it’s possible that some questions in our survey were ambiguous, leading to variable responses from participants. For example, when asked to name a plot, some participants tended to describe it. We might have gotten more consistent results by asking people to name plots in two or one words, or by asking: “What would you call a software function to plot this figure? Similarly, some of our questions about software features might be ambiguous, for example there is some room for interpretation in how to understand ‘interactivity’ or ‘reproducibility’.

Finally, our study only covers plots of event-related potentials. It remains to be seen which findings can be generalized to other EEG plots in the domains of spectral estimation or connectivity (De Ridder et al., 2015, 2019).

### 5.10. Further research

A survey like ours only captures a still image of an ongoing process. The field is self-educating, software is constantly improving, or new visualizations are being proposed. For future studies on this topic, we suggest some interesting follow-up questions:

- In addition to our question of which tool the respondents have experience with, one could ask “What is the main tool you use for ERP analysis?” to find out which tool is in use most of the time.
- “If multiple plot types are available to display similar information, how do you choose which to use (e.g. topoplots vs. butterfly plot vs. channel image)?
- “If you did not use all electrodes in your research, what was the reason for this decision?”
- “Are you familiar with sequential and diverging color maps for data visualization?”
- “What is your default color map?”

Furthermore, an interesting approach for future studies might be to present participants with figures that follow or violate certain visualization principles and ask participants to identify potential visualization problems or shortcomings. This method would allow for a more indirect assessment of awareness of visualization problems.

We found it noteworthy that no study to date has investigated how interpolation algorithms, interpolation shape (circle, neck, elongated), or projection algorithms affect the perceived strength of evidence and implicit localization of EEG activity on topoplots. The difference in accuracy and precision between plotting time points and averaged time periods in topoplot series is not studied either. In addition, we are not aware of a study investigating the length of the baseline period and its influence on being able to gauge the signal-to-noise ratio.

A separate topic could be a study of EEG visualization practices in research institutions in Asia, Africa, and South America, and in medical or practical settings since such respondents were lacking in our study. It will also be interesting to repeat a similar study in ten years to see what progress has been made in the field.

## 6. Conclusions

This study provides insight into the current state of ERP visualization in the EEG community. In our study we identified the most popular tools among EEG researchers, understood preferred features of visualization tools, addressed common opinions on controversial topics, and assessed visualization literacy. We also identified areas for improvement in EEG analysis tools and suggested some topics for future research. The results of this study can guide the development of more effective and user-friendly ERP visualization tools, ultimately advancing the field of EEG research and facilitating more accurate and interpretable data analysis.

## 7. Ending sections

### 7.1. Data and code availability

The data and analysis script are publicly available. The raw dataset can be found at the DaRUS-Dataverse repository https://doi.org/10.18419/darus-3729 (Mikheev, Skukies, et al., 2023a). Analysis scripts are available at https://github.com/vladdez/Survey_analyses/tree/main or https://zenodo.org/records/10402375 and can be cited as (Mikheev, Skukies, et al., 2023b).

### 7.2. Author Contributions

**Vladimir Mikheev:** Conceptualization, Software, Methodology, Data Curation, Investigation, Writing - Original Draft, Visualization, Formal analysis.

**René Skukies**: Methodology, Writing - Review & Editing.

**Benedikt Ehinger**: Conceptualization, Methodology, Writing - Review & Editing, Supervision, Visualization.

### 7.3. Funding

Funded by the Deutsche Forschungsgemeinschaft (DFG, German Research Foundation) – Project-ID 251654672 – TRR 161. Additionally, René Skukies was funded by Deutsche Forschungsgemeinschaft (DFG, German Research Foundation) under Germany’s Excellence Strategy - EXC 2075 - 390740016.

## 7.5. Acknowledgments

We would also like to acknowledge Judith Schepers, Jan Haas, Martin Geiger, Hannes Bonasch, Guanhua Zhang, Luis Lips, Felix Schroder, Olaf Dimigen, Robert Oostenveld, Phillip Alday for their feedback on the methodology, structure, and design of the survey.

## Supplementary materials

**Figure 1.**
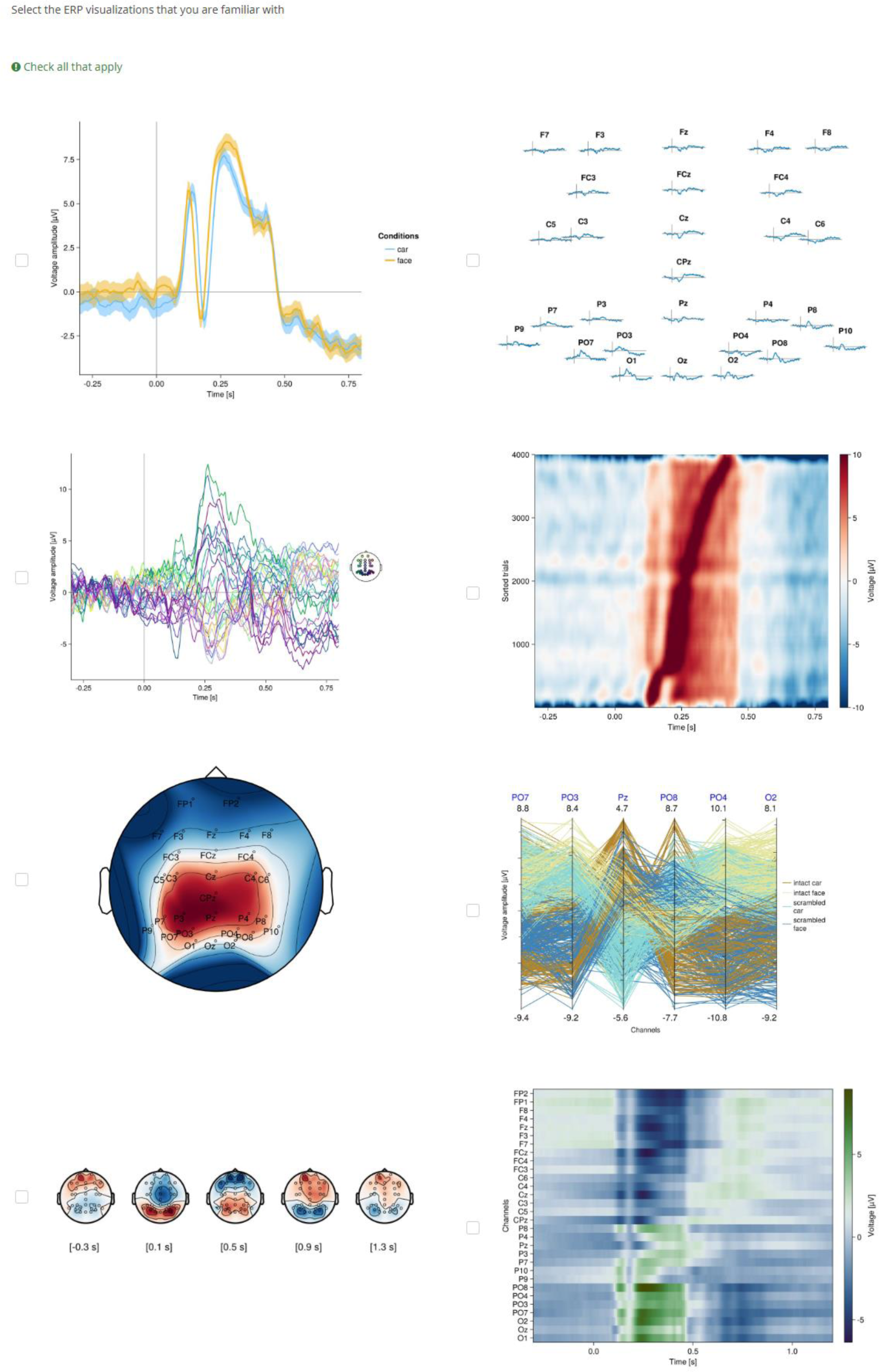
These plots were shown in the survey. Participants indicated which they recognize.

**Figure 2.**
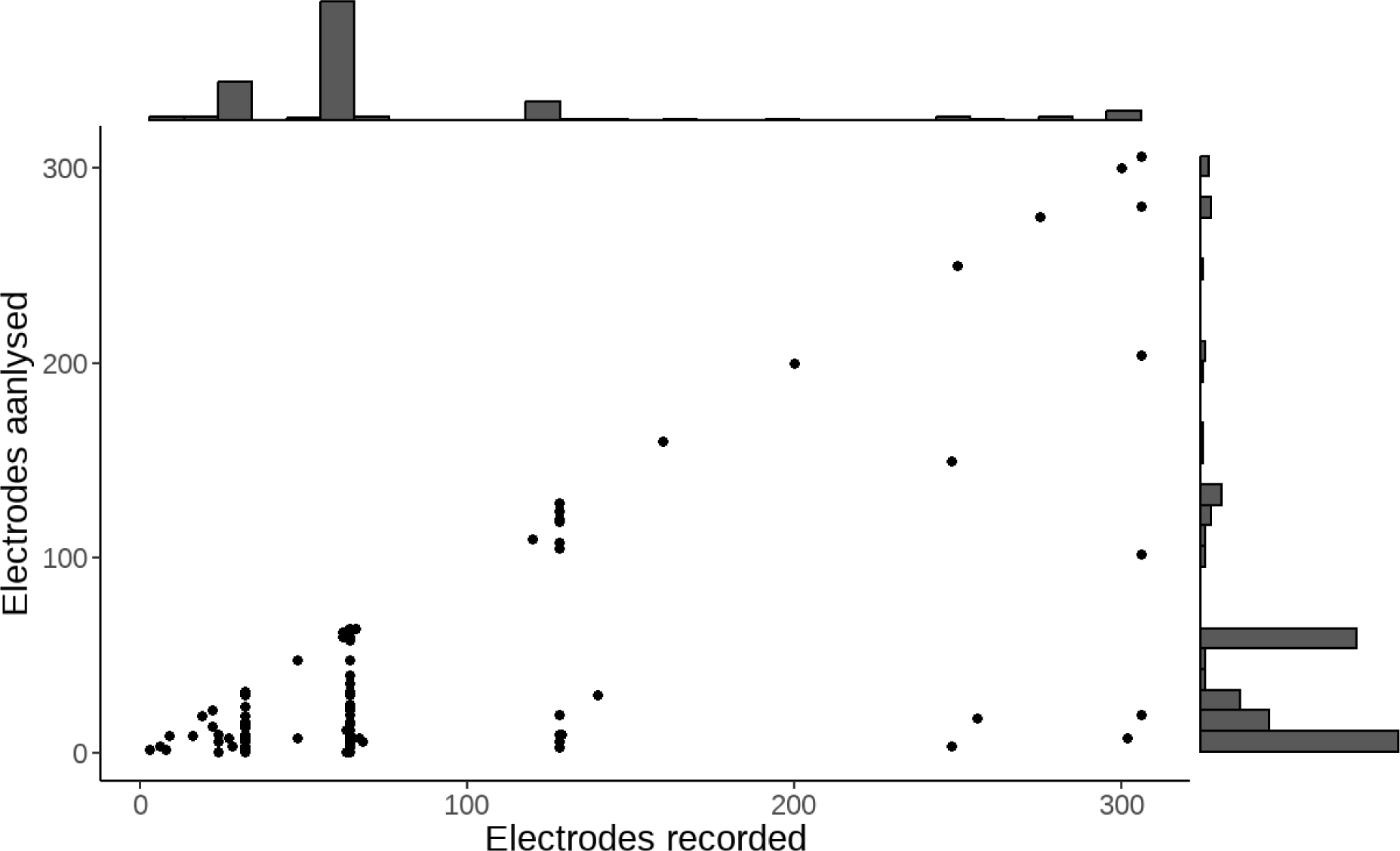
A marginal histogram of number of electrodes analyzed and recorded (N of respondents - 208)

**Table 1.**
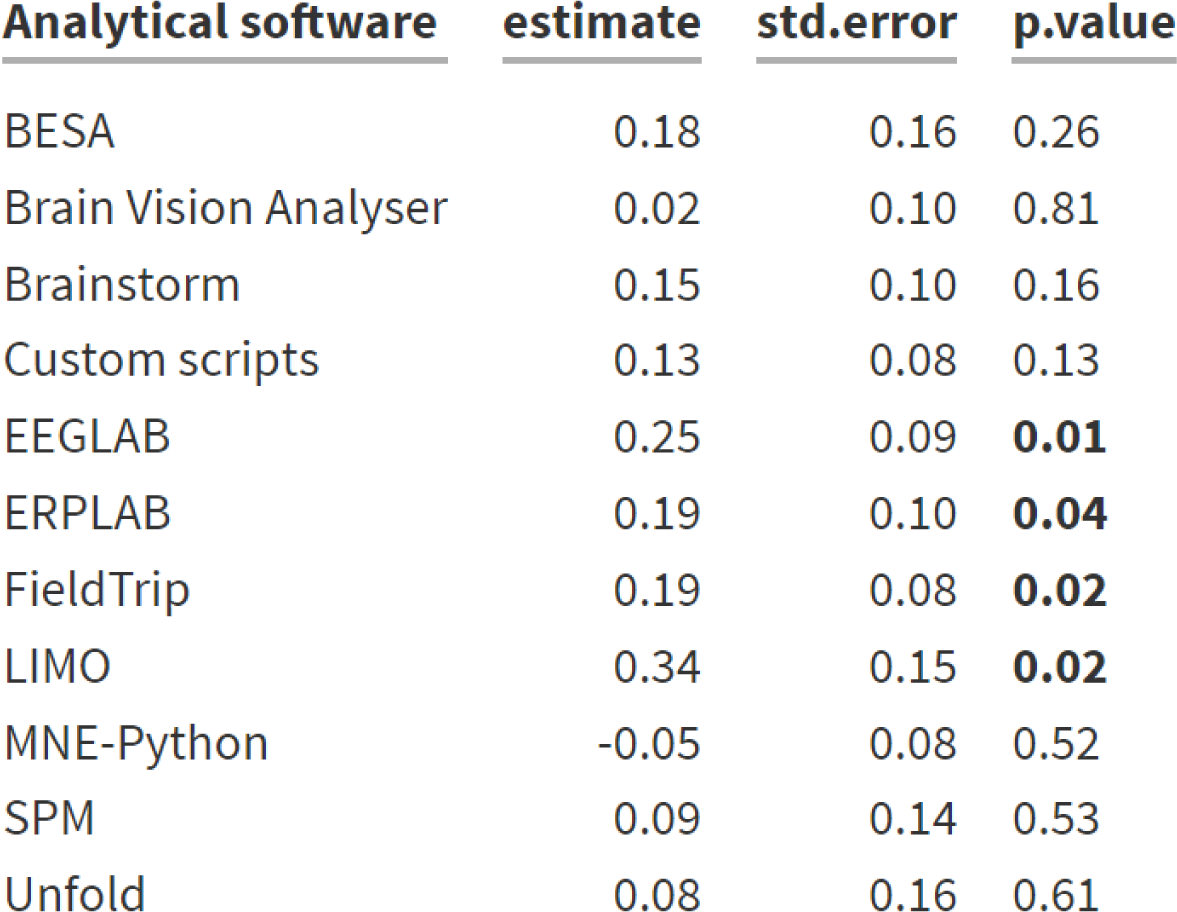
Results of logistic regressions where the dependent variables are the analytical tools, and the independent variable is the researcher’s proficiency (N of respondents - 210)

| <u>Analytical software</u> | <u>estimate</u> | <u>std.error</u> | <u>p.value</u> |
| --- | --- | --- | --- |
| BESA | 0.18 | 0.16 | 0.26 |
| Brain Vision Analyser | 0.02 | 0.10 | 0.81 |
| Brainstorm | 0.15 | 0.10 | 0.16 |
| Custom scripts | 0.13 | 0.08 | 0.13 |
| EEGLAB | 0.25 | 0.09 | **0.01** |
| ERPLAB | 0.19 | 0.10 | **0.04** |
| FieldTrip | 0.19 | 0.08 | **0.02** |
| LIMO | 0.34 | 0.15 | **0.02** |
| MNE-Python | -0.05 | 0.08 | 0.52 |
| SPM | 0.09 | 0.14 | 0.53 |
| Unfold | 0.08 | 0.16 | 0.61 |

**Table 3.**
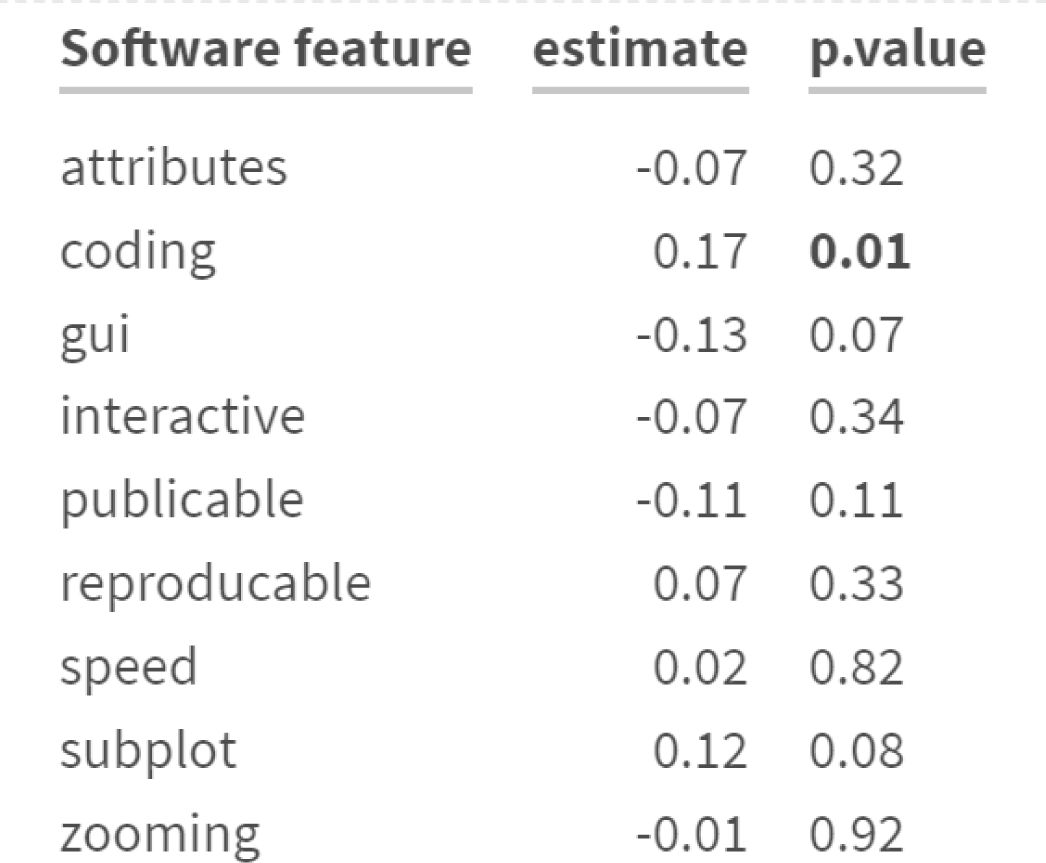
Results of Spearman’s rank correlation tests where the dependent variables are the features of analytical tools, and the independent variable is the researcher’s proficiency (N of respondents - 210)

## Footnotes

1 Channel plot - a heatmap similar to a single-trial ERP image, but depicting time and channel activity, which is commonly used in e.g. the LIMO toolbox (Pernet et al., 2011)

2 One of the reasons for this is that the paper is currently not widely cited (36 citations on Google Scholar, none from EEG studies), most likely due to a mismatch between the communities of visualization researchers and ERP practitioners.

3 Early pioneers of electroencephalography, such as Richard Caton, Hans Berger, Alfred Lumis, and Edgar Adrian, did not consistently indicate polarity notation (Adrian & Matthews, 1934; Berger, 1929; Caton, 1875; Loomis et al., 1935). In one of the earliest cases we could find, where the sign of EEG waveforms was noted (Prawdicz-Neminski, 1925, dog EEG) the positive was up, but the time axis was reversed from right to left, which is of course even more contrary to current standards.

4 The respective comparison data are not part of the original paper but can be found at https://neuro.debian.net/survey/2011/results.html

5 Only 49% of them used RGB in full spectrum and in a standard order of colors.

6 34% of the papers had at least one figure with a rainbow color map and 21% contained a figure with red-green elements.

7 34 papers used the EK500 color map, which is even worse, but was the default color scheme on the Simrad EK500, one of the first scientific echosounders. It has only 12 available colors due to the limited capabilities of a bit plane, although echosounders have become more advanced since then.

